# Intermediate, rather than Isohydric or Anisohydric, Species were More Resilient to an Extreme Heatwave in a Common Garden in Sydney

**DOI:** 10.64898/2026.10.23.761478

**Authors:** Wahidullah Rahmani, Mohammad Golam Kibria, Mark G. Tjoelker, Paul D. Rymer, Renée M. Marchin

## Abstract

Urban forests are increasingly exposed to climate extremes, yet selecting resilient urban species remains difficult as it is unclear why some species fail and others succeed. We monitored growth and survival of 48 urban tree and shrub species, respectively, for about five years after planting, plus we quantified species’ crown dieback after a natural extreme heatwave (three days >40 °C) in a common garden in Sydney, Australia. For 30 species we measured 15 plant functional traits, including both hydraulic and morphological traits. Native species had less crown dieback and higher absolute growth but lower survival than exotic species. Plant functional traits linked to heat and drought tolerance explained 20-30% of the variation in observed crown dieback across species. Extensive dieback >40% after the extreme heatwave was found for seven species with large leaf size (and leaf width), high leaf relative water content, and low wood density. Nonlinear relationships between crown dieback and leaf water potential traits identified that both isohydric and anisohydric species were vulnerable, indicating that vulnerability did not align neatly with a single hydraulic strategy. Hydraulic failure likely occurred for some, but not all, vulnerable isohydric and anisohydric species, suggesting vulnerability of certain species was directly caused by cumulative thermal stress. Importantly, survival was not related to any single functional trait, indicating that crown dieback was a more sensitive and earlier indicator of cumulative thermal stress than mortality. These findings support a shift from climate of origin-based to trait-based species selection in urban forestry.

**Highlight:**

- Crown dieback was an earlier indicator of cumulative thermal stress than mortality.
- Functional traits explained 20–30% of variation in crown dieback among 30 species.
- Both isohydric and anisohydric species were vulnerable to extreme heat.
- Trait-based selection may improve climate resilience of urban forests.

## Introduction

Healthy and resilient urban forests are fundamental to the long-term liveability of cities but are threatened by climate change (Esperon-Rodriguez et al., 2022; Zhu et al., 2025). The increasing severity and frequency of extreme events (e.g., droughts, heatwaves, storms, wildfires) disrupts physiological function, constrains growth, and increases the incidence of crown dieback and mortality of urban trees (Marchin et al., 2022a; Brandt et al., 2016; Smith et al., 2019). As a result, substantial declines in urban tree survival are projected globally (Burley et al., 2019; Esperon-Rodriguez et al., 2019; Sjöman et al., 2024). These threats are already evident, with recent extreme heat events causing widespread leaf loss across urban canopies (Franceschi et al., 2023; Marchin et al., 2022b; Tabassum et al., 2021; Sanusi and Livesley, 2020). Heatwaves may be particularly hazardous for urban trees due to the urban heat island (UHI) effect, a commonly observed phenomenon characterised by higher air temperatures (∼4-5 °C) in urban areas compared to neighbouring suburban and rural environments (Oke, 1982; Santamouris, 2020). This effect is primarily driven by urban structures, including heat-absorbing materials like asphalt and concrete, but also by limited vegetation and anthropogenic heat emissions in cities (Cuce et al., 2025). Urban trees are essential city infrastructure (Esperon-Rodriguez et al., 2026) that help mitigate the UHI and reduce local air and surface temperatures by cooling the environment through transpiration (Winbourne et al., 2020), shading buildings and paved surfaces to reduce energy demand (Wang et al., 2018), and sequestering carbon in long-lived biomass (Nowak & Crane, 2002; Petri et al., 2016; Pregitzer et al., 2022; Zhao et al., 2023). Consequently, urban practitioners must prioritise the selection of heat-tolerant species to avoid poor returns on investments in planting and maintenance, plus the loss of ecosystem services, such as urban cooling, during hot summer periods. Accordingly, identifying and establishing tree species resilient to future climate conditions has become a central goal of urban forest planning (Brandt et al., 2016).

Despite a growing recognition of climate risks to urban forests, identification of climate-resilient species remains difficult in part because optimal species vary with climate and by city (Chen et al., 2024; Marchin et al., 2025), but also due to the need to balance tree species with available water (Huang et al., 2020). The heat tolerance of tree species depends on a combination of functional traits that dictate the leaf energy budget, including leaf width, thickness, relative water content, and stomatal conductance (Leigh et al., 2012; Michaletz et al., 2016) These traits are also known to vary with species’ drought strategy, which can be characterised along an isohydric to anisohydric continuum, where isohydric species close stomata early during drought to maintain high (less negative) leaf water potentials, while anisohydric species generally sustain gas exchange for longer, allowing leaf water potential to decline (Tardieu & Simonneau, 1998; Klein, 2014). The leaf water potential at turgor loss point (π_tlp_) is one of the most widely used leaf traits for assessing drought tolerance across species, as it indicates the water potential at which leaf cells lose turgor and leaves wilt (Bartlett et al., 2016; Maréchaux et al., 2015; Sjöman et al., 2018). Species with low π_tlp_ have higher leaf-level drought tolerance, resulting in their higher survival in common gardens in both a temperate and subtropical city in Australia (Kibria et al., 2026). The π_tlp_ has also been used to indicate species’ degree of isohydry, with high π_tlp_ characteristic of isohydric species (Fu and Meinzer, 2019). Fast-growing isohydric species with high π_tlp_, low wood density, and large, thin leaves experienced greater crown dieback and mortality during Sydney’s extreme 2019–2020 summer (Marchin et al., 2022a), supporting the commonly observed growth– survival trade-off under stressful conditions (Greenwood et al., 2017; O’Brien et al., 2017). Interestingly, drought-tolerant species with low π_tlp_ can still achieve higher growth in cities, relative to species with higher π_tlp_ (Kibria et al., 2024), suggesting π_tlp_ is a key trait for selection of urban species.

Effective urban forest management requires integrating climate-resilient species selection with evidence derived from long-term monitoring of tree performance, survival, and crown dieback, particularly to identify species that can withstand cumulative thermal stress and repeated exposure to extreme heat. Quantifying the risk of species decline and mortality is therefore essential for urban forest managers seeking to optimise resource allocation and minimise losses in planting programs (Esperon-Rodriguez et al., 2022; Hilbert et al., 2019). Common-garden plantings established under operational urban-forestry conditions provide standardised, field-based comparisons of growth, early decline, and mortality across tree species. For example, an Australian urban trial network using standardised plantings across multiple cities reported substantial early mortality of 23% of planted individuals and found strong site effects (Esperon-Rodriguez et al., 2025b). Sites on dense urban streets often have higher growth and mortality than those in parks and gardens (Esperon-Rodriguez et al., 2025b; Esperon-Rodriguez et al., 2023), although park trees experienced greater water stress than street trees in a dry Mediterranean city (Dekel et al., 2026). Heatwaves are intensifying and now occur regularly in cities, so there is a both a need and opportunity to identify the long-term effects of heat stress on urban trees (Marchin et al., 2025), including which species are more at risk.

Strategic urban greening strategies must account for the slow growth rates of trees and the importance of promoting longevity, requiring planning decades in advance and selecting urban tree species that will be tolerant of future climate changes (Esperon-Rodriguez et al., 2024a). To address this need, a common garden experiment was established in western Sydney, Australia, as it represents one of the city’s hottest suburbs, providing a valuable lens into plant performance under future warmer climates. A previous study by Kibria et al. (2026) used this common garden to assess survival during the two-year establishment phase, quantifying early growth and mortality across a diverse planting of 48 tree and shrub species. Building on this foundation, we continued measurements into the post-establishment phase (i.e., up to about five years after planting) to assess longer-term growth, early indicators of decline, and survival across a transition from relatively favourable establishment conditions to a hotter and drier climate during post-establishment. For a subset of 30 species, we also measured 15 plant functional traits related to plant water relations, drought tolerance, and/or heat tolerance, along with their crown dieback following a three-day extreme heatwave (>40 °C). First, we determined how patterns of growth and mortality changed over time (i.e., from the first two establishment years to the later post-establishment phase) before addressing the following questions: (1) Did growth, crown dieback, and mortality differ between native Australian and exotic species? (2) Which plant functional traits predicted crown dieback and survival across species? We hypothesised that native Australian species would have higher growth and survival, plus lower crown dieback after the extreme heatwave, relative to exotic species, reflecting adaptation to local climate conditions. We also expected that species with more conservative, anisohydric functional traits, such as smaller leaves, higher leaf mass per area, higher wood density, higher stomatal conductance under high atmospheric vapour pressure deficit, and lower relative water content during hot periods were expected to have higher survival and lower crown dieback than more isohydric species after cumulative heat stress. Our findings provide critical information for the selection of urban tree species for the development of resilient green cities under climate change.

## Materials and Methodology

### Study Site and Species

An experimental common garden was established in Richmond (33°37′15′′S, 150°45′15′′E) in western Sydney, New South Wales (NSW), Australia on the university campus at the Hawkesbury Institute for the Environment of Western Sydney University (Fig. 1). The long-term climate in Richmond is subtropical characterized by hot summers with a mean annual temperature (MAT) of 17.8°C and mean annual precipitation (MAP) of 765 mm. Based on long-term climate records (1993-2025), January is the hottest month with an average temperature of 30.4°C, while July is the coldest and driest month at 17.9°C and 35.7 mm of rainfall (Fig. 2). February is the wettest month, receiving an average of 113.0 mm of rainfall. Climate data is from a Bureau of Meteorology station (BoM 067105, Australian Bureau of Statistics, 2025) located 3.5 km from the study site. The experimental site was planted in December 2020 and covers an area of 150 m by 185 m, or 2.8 ha. The study site contains 48 species, including 34 native Australian and 14 exotic species (Table 1, Table S1), with 32 plants per species distributed across four blocks, resulting in a total of 1,536 plants. Species were selected from a wide range of climate origins, from warm to cool, wet to dry, and narrow to broad climatic origins (Table 1; for further details, see Kibria et al., 2026).

**Figure 1:**
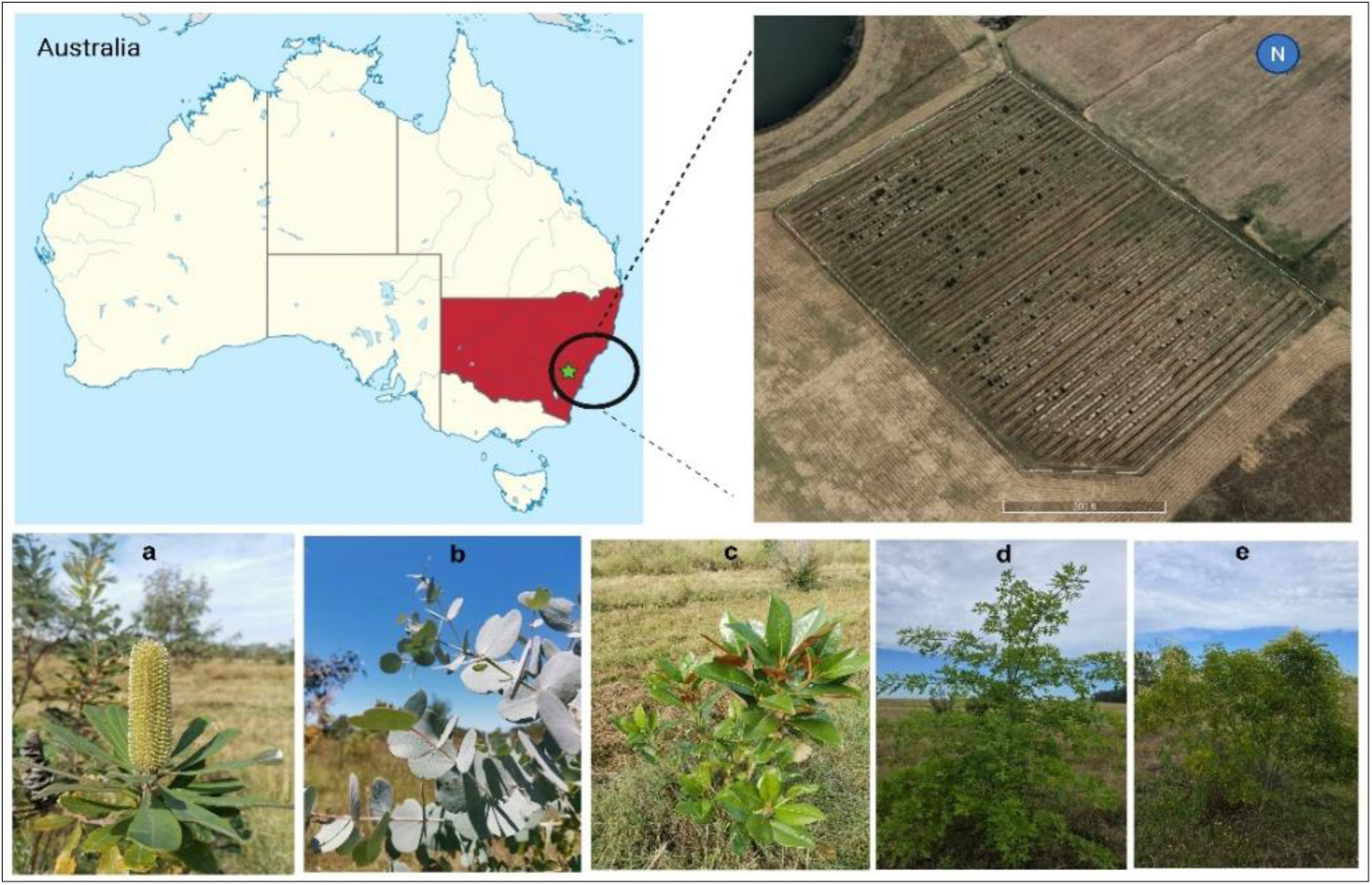
Map of study site located in Richmond, NSW, Australia, showing the experimental layout and examples of study species, including (a) Banksia integrifolia, (b) Eucalyptus rubida, (c) Magnolia grandiflora, (d) Quercus palustris, and (e) Fraxinus griffithii.

**Figure 2.**
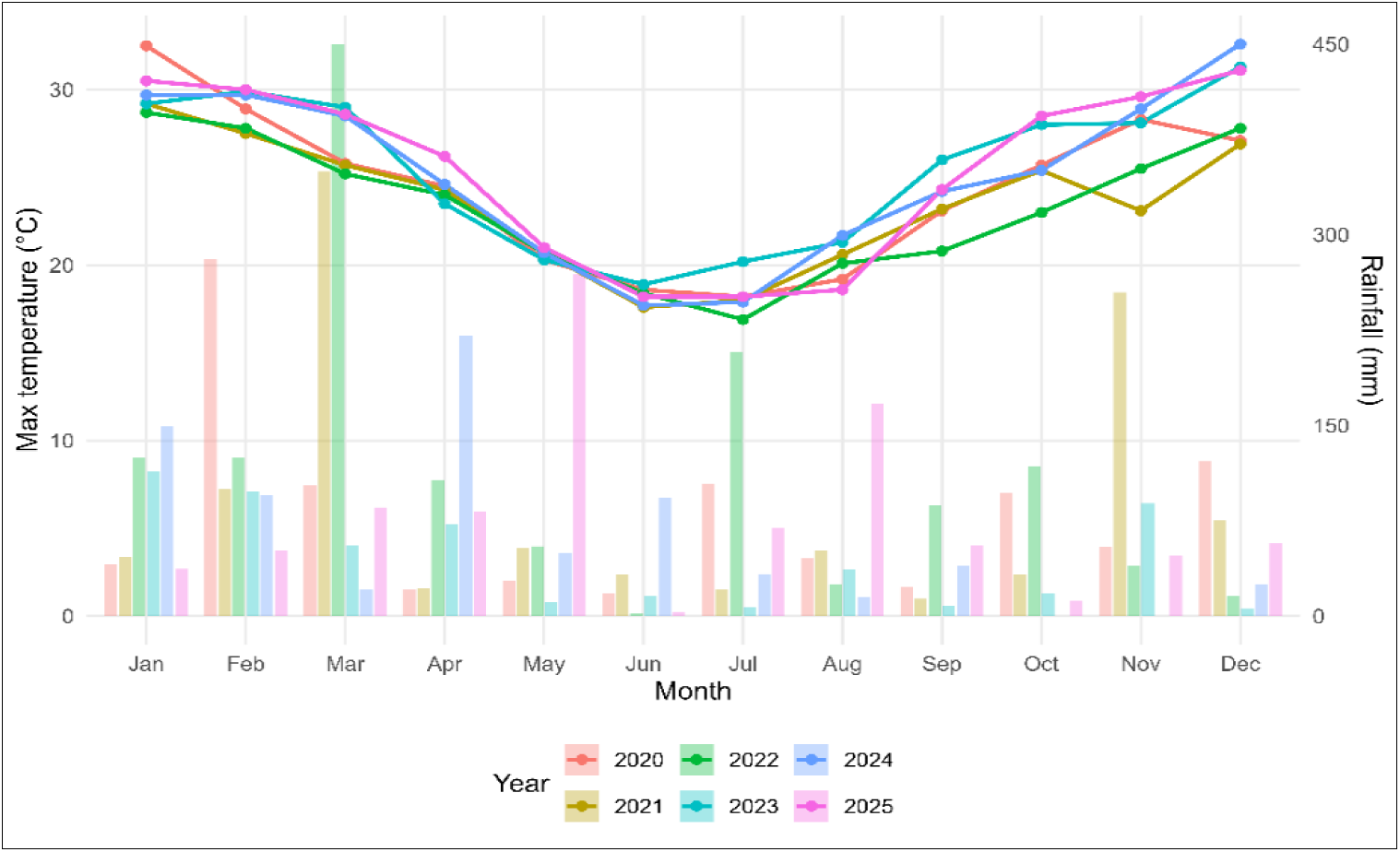
Seasonal climate patterns during the study (2020-2025). Points show mean monthly maximum temperature (°C), while bars represent total monthly rainfall (mm). Climate data provided by the Bureau of Meteorology station 067105.

**Table 1:** Thirty species-level hydraulic traits used to derive degree of isohydry continuum. Values are means **±** SE for leaf water potential at turgor loss point (π_tlp_, MPa), wood density (g cm⁻³), and daily peak leaf water potential decline (ΔΨ, MPa), or the largest ΔΨ value from four monthly samplings throughout the 2024-2025 austral summer. MAT (mean annual temperature, MAP (mean annual precipitation). Species are ordered by degree of isohydry from most isohydric to most anisohydric.

| Species | MAT | MAP | Iso/anisohydric | Wood density | $\pi_{\text{tlp}}$ (MPa) | $\Delta\Psi$ (MPa) |
| --- | --- | --- | --- | --- | --- | --- |
| *Syzygium luehmannii* | 19.100 | 1629.000 | Isohydric | 0.477± 0.008 | -1.618 ± 0.047 | -0.090 ± 0.067 |
| *Acacia implexa* | 15.200 | 784.700 | Isohydric | 0.589 ± 0.067 | -1.646 ± 0.181 | -0.245 ± 0.390 |
| *Elaeocarpus reticulatus* | 17.400 | 1256.200 | Isohydric | 0.434 ± 0.022 | -2.085 ± 0.046 | -0.157 ± 0.050 |
| *Eucalyptus mannifera* | 12.500 | 819.400 | Isohydric | 0.583 ± 0.028 | -1.606 ± 0.027 | -0.620 ± 0.142 |
| *Eucalyptus rubida* | 13.200 | 929.000 | | 0.514 ± 0.014 | -1.546 ± 0.042 | -1.407 ± 0.089 |
| *Fraxinus griffithii* | 20.500 | 2170.200 | | 0.585 ± 0.016 | -1.736 ± 0.032 | -0.462 ± 0.191 |
| *Tristaniopsis laurina* | 16.100 | 1096.200 | | 0.555 ± 0.014 | -1.974 ± 0.061 | -0.107 ± 0.043 |
| *Banksia integrifolia* | 16.400 | 1235.700 | | 0.548 ± 0.099 | -1.814 ± 0.086 | -0.545 ± 0.231 |
| *Corymbia eximia* | 16.700 | 993.300 | | 0.536 ± 0.011 | -1.691 ± 0.059 | -1.180 ± 0.132 |
| *Magnolia grandiflora* | 18.200 | 1320.500 | | 0.375 ± 0.029 | -1.75 ± 0.027 | -1.372 ± 0.051 |
| *Eucalyptus globulus* | 14.900 | 777.500 | | 0.507 ± 0.038 | -1.78 ± 0.062 | -0.975 ± 0.129 |
| *Flindersia australis* | 19.400 | 1122.800 | | 0.604 ± 0.012 | -1.975 ± 0.054 | -0.280 ± 0.106 |
| *Liquidamber styraciflua* | 15.800 | 1216.000 | | 0.538 ± 0.018 | -1.964 ± 0.076 | -0.415 ± 0.078 |
| *Lophostemon confertus* | 19.000 | 1316.500 | Mesohydric | 0.510 ± 0.008 | -1.782 ± 0.049 | -0.565 ± 0.385 |
| *Syncarpia glomulifera* | 17.600 | 1229.900 | Mesohydric | 0.732 ± 0.047 | -1.677 ± 0.023 | -0.312 ± 0.062 |
| *Ulmus parvifolia* | 17.700 | 1130.800 | Mesohydric | 0.646 ± 0.067 | -1.75 ± 0.036 | -0.525 ± 0.027 |
| *Acacia cognata* | 14.400 | 926.400 | Mesohydric | 0.698 ± 0.038 | -1.66 ± 0.099 | -0.537 ± 0.161 |
| *Acacia longifolia* | 15.600 | 831.300 | | 0.666 ± 0.035 | -1.54 ± 0.055 | -1.632 ± 0.155 |
| *Cupaniopsis anacardioides* | 20.700 | 1226.100 | | 0.808 ± 0.062 | -1.75 ± 0.059 | -0.445 ± 0.100 |
| *Eucalyptus bancroftii* | 18.900 | 1247.400 | | 0.614 ± 0.019 | -1.76 ± 0.060 | -0.677 ± 0.084 |
| *Eucalyptus pauciflora* | 9.600 | 1142.900 | | 0.661± 0.071 | -1.656± 0.009 | -0.692 ± 0.022 |
| *Acacia melanoxylon* | 14.400 | 969.100 | | 0.631 ± 0.011 | -1.589 ± 0.039 | -1.110 ± 0.085 |
| *Quercus ilex* | 13.000 | 740.600 | | 0.718 ± 0.012 | -2.08 ± 0.249 | -0.375 ± 0.104 |
| *Alphitonia excelsa* | 19.600 | 1183.100 | | 0.465 ± 0.012 | -2.31 ± 0.213 | -1.712 ± 0.237 |
| *Fraxinus oxycarpa* | 11.300 | 679.700 | | 0.659 ± 0.030 | -1.93 ± 0.089 | -1.382 ± 0.128 |
| *Quercus dentata* | 10.900 | 1254.100 | | 0.634 ± 0.027 | -2.20 ± 0.121 | -0.977 ± 0.308 |
| *Callistemon viminalis* | 17.700 | 1056.000 | Anisohydric | 0.672 ± 0.013 | -1.75 ± 0.100 | -1.965 ± 0.254 |
| *Callistemon pallidus* | 12.600 | 1016.900 | Anisohydric | 0.686 ± 0.013 | -2.09 ± 0.152 | -1.085 ± 0.041 |
| *Quercus palustris* | 11.300 | 947.100 | Anisohydric | 0.660 ± 0.020 | -1.98 ± 0.073 | -2.242 ± 0.629 |
| *Pyrus nivalis* | 9.300 | 920.200 | Anisohydric | 0.636 ± 0.018 | -2.48 ± 0.063 | -2.705 ± 0.202 |

### Species Growth, Dieback, and Mortality

Mortality, tree height (ground level to top of stem; H, cm) and basal diameter (stem diameter at 10 cm above the ground; D, mm) were recorded for all 1,536 trees (i.e., 48 species, 32 trees per species) annually in December during the establishment phase (i.e., December 2021, 2022; see Kibria et al. (2026) for this previously reported data), then annually at the end of the growing season during the post-establishment phase (i.e., April 2024, May 2025) at 3.5 and 4.5 years, respectively, after planting. Absolute growth rates (AGR) and relative growth rates (RGR) were calculated from tree height and diameter measurements for each species and year. There were 10 species with fewer than four individuals in 2025 (Table S1) that were excluded from height and diameter growth analyses. Applying this threshold resulted in 38 species being included in height and diameter growth analyses. AGR (cm·year⁻¹) was calculated as: AGR = (H̅_2025_ - H̅_2021_)/ (t_2_ - t_1_), and RGR (cm·cm⁻¹·yr⁻¹) was estimated as: RGR = (ln[H_t2_ or D_t2_] - ln[H_t1_ or D_t1_])/(t_2_-t_1_), where t_1_ and t_2_ are the times of the first (in 2021) and last (in 2025) measurement, respectively.

Here, mortality is defined as the irreversible loss of physiological function, typically indicated by the complete absence of live green foliage outside the normal leaf-off period (for deciduous species), accompanied by evidence of whole-tree death, such as the absence of live buds or expanding leaves, presence of dead stems, lack of re-sprouting, and failure to respond to favourable growth conditions. Annual mortality was calculated from the number of apparently dead individuals each year. Species mean mortality was calculated as the percentage of leafless trees divided by the total number of live trees in the previous sampling date (Mortality 2021 = leafless in 2021 / live in 2020; Mortality 2022 = leafless in 2022 / live in 2021; Mortality 2024 = leafless in 2024 / live in 2022; Mortality 2025 = leafless in 2025 / live in 2024) and as cumulative mortality (Mortality 2025 = leafless in 2025 / planted in 2020).

Crown dieback was assessed after a naturally occurring three-day extreme heatwave (8-10 January 2026, maximum T_air_ = 41.9, 42.0, 42.8 °C) using a visual severity index scored from 0 to 100 (0 = no dieback; 100 = complete defoliation or leaf death). Crown dieback was quantified as the proportion of each tree crown showing visible stress symptoms, including leaf burn, desiccation, or premature leaf drop, relative to a healthy individual with the typical crown form for that species. A total of 32 trees per species were assessed, and each plant was scored independently by two assessors from multiple angles, with observer scores averaged prior to analysis.

### Species’ Drought Strategy

Species were ranked along the isohydric to anisohydric continuum by integrating data from three plant functional traits related to species’ drought strategy: (1) leaf water potential at turgor loss point (π_tlp_), (2) wood density, and (3) daily leaf water potential decline (ΔΨ; Table 1). For each trait, species were ordered and then assigned a rank from 1 (most isohydric) to 20 or 24 (most anisohydric); the same rank was given to species with similar trait values (i.e., < mean standard error). Ranks from all three traits were summed for the final classification, and the four lowest rankings were ‘isohydric’, the four intermediate rankings were ‘mesohydric’, and the four highest rankings were ‘anisohydric’. Leaf water potential at turgor loss point (π_tlp_) was estimated by measuring leaf osmotic potential (π_o_; Bartlett et al. 2012) for 21 species in December 2025 and for 9 species pooled across two sampling years (2022, 2025; Kibria et al., 2026). Healthy and fully-expanded leaves were collected from each species (n=4 trees per species) and rehydrated in distilled water overnight (12-14 h; Barros et al., 2022). We removed the midrib from all fully-hydrated leaves (i.e., leaf water potential > -0.3 MPa) and 40-mm diameter leaf discs or leaf pieces were quickly cut from each leaf. Leaf discs were wrapped in foil and frozen in liquid N^2^ for two minutes and then equilibrated for 10 min inside a sealed, humidified plastic bag. Leaf sections were punctured repeatedly with sharp-tipped forceps before measurement. We used an osmometer (WP4C Dewpoint PotentiaMeter, Decagon Devices, Pullman, WA, USA) and recorded measurements for 30-42 min, until equilibrium as indicated by <0.01 MPa change over two minutes. Finally, π_tlp_ was calculated from osmotic potential (π_o_) using the equation: π_tlp_ = 0.832 π_o_ -0.631 (Bartlett et al., 2012). Wood density (n = 4 trees per species) was determined in December 2025 for an 8-10 cm stem segment after splitting to remove the pith and bark. Fresh sapwood volume was determined using the water displacement method before wood samples were oven dried to a constant mass at 105°C (Slick et al., 2007).

Leaf water potentials were measured on leaves (n= 4 trees per species) using a pressure chamber (Model 1505D, PMS Instruments, Corvallis, OR, USA) for all 30 species on four different days (December 2024, January, February, March 2025) throughout the austral 2024-2025 summer, selecting hot days (maximum T_air_ = 36.9, 39.2, 42.5, 31.3°C). To measure predawn leaf water potential (Ψ_PD_), healthy and fully-expanded leaves were randomly sampled from the north side of each tree (n=1 leaf per tree) at the early morning (5:00-6:00), kept in sealed plastic bags containing a small piece of wet paper towel, and stored in cool, dark conditions prior to measurement. For midday leaf water potential (Ψ_MD_), one leaf per tree was sampled in the middle of the day (12:30-13:30) on the same day as Ψ_PD_, following the same procedure. All measurements were completed within 4 h for each predawn and midday collection. A dissecting microscope was placed above the pressure chamber, with fibre-optic lights, to clearly observe the balance pressure, which was recorded as the mid-point between the first sign of water from the petiole and when the entire xylem area was wetted (Rodriguez-Dominguez et al., 2022). All repeated Ψ_PD_ and Ψ_MD_ measurements were averaged across individuals and species for analysis. The largest peak ΔΨ for each species was selected from the four sampling dates after calculation using the equation: Ψ_MD_ − Ψ_PD_.

### Plant Morphological Traits

Functional traits were collected for a subset of 30 species, primarily during the 2024-2025 austral summer from December 2024 to January 2025. For leaf size, leaf mass area (LMA), leaf thickness and leaf width, ten healthy and fully-expanded leaves were sampled from the north side of four tree per species and kept in sealed plastic, humidified bags and stored in cool, dark conditions prior to measurement. Leaf size of fresh leaves (n=10 leaves per species) was measured using a flatbed scanner WinFOLIA Pro 2005 software (Regent Instruments Inc.), from which leaf width (cm) and leaf area (cm^2^) were measured. LMA was calculated as the ratio of the leaf surface area to dry mass (cm^2^ g^-1^). Leaf thickness (mm) was measured on the same leaves using digital callipers, taking three readings per leaf (avoiding the midrib) and using the mean for analysis. Relative water content (RWC) was measured using the excised leaf method (McCaig & Romagosa, 1989) and calculated as: (FM - DM)/(TM - DM) × 100 (Wang et al., 2019). Freshly excised leaves (n= 10 leaves per species) were weighed for fresh mass (FM), rehydrated to full turgor in the dark to obtain turgid mass (TM), then oven-dried at 70°C for 48 h to constant mass for dry mass (DM).

Stomatal density was determined using the stomatal imprint method (Rahman et al., 2022). Nail polish stomatal imprints were successfully obtained for all 30 species using leaves collected in December 2024 to January 2025. Stomatal imprints were collected from the abaxial surface of leaves. Midribs were avoided while taking the imprints. The area halfway between the midrib and margin of the leaf and ∼1/3 down toward the petiole end was painted using transparent quick dry nail polish to standardise the stomatal imprint collection. The imprints were then extracted using a transparent sticky tape (approximately 10 x 18 mm size of the impression peel) once the nail polish samples were dried and adhered to microscope slides. A camera mounted microscope (Zeiss Axio Scope A1, Germany) was used to observe and capture the images of stomata for the determination of stomata density in Image-J software (Schneider et al., 2012). Three images using three different fields of view per microscopic slide were taken using 20x magnification. Stomatal density (SD, the number of stomata per mm) on the leaf abaxial surface was determined from each image and averaged for one leaf sample using 20x magnification image (Yan et al., 2012).

### Leaf temperature and gas exchange

Maximum leaf temperature (T_leaf,max_) was measured on sun-exposed leaves (n = 4 trees per species) on the last day of the three-day extreme heatwave (10 January 2026, maximum T_air_ = 42.8 °C). Measurements were taken at the hottest time of day (13:00-15:00) using an infrared thermal camera (T640, FLIR Systems, Wilsonville, OR, USA) to capture peak leaf surface temperature. Stomatal conductance (*g_s_*), leaf temperature, and effective quantum yield of photosystem II (Φ_PSII_) were measured using a LI-600 Porometer/Fluorometer (LI-COR Biosciences, Lincoln, Nebraska, USA) on healthy, fully-expanded leaves at midday (13:00-14:00) on six hot days (>35°C) from December 2024 to April 2025 for all 30 species (n=4 trees per species). All repeated g_s_, leaf temperature, and Φ_PSII_ measurements were averaged across individuals and species for analysis.

### Statistical Analyses

We tested if growth and mortality varied across species and over time using full-factorial, two-way analyses of variance (ANOVA) with species, year, and their interaction as fixed effects. Survival was additionally modelled using a binomial generalized linear model (GLM) fitted to counts of alive individuals with year as a fixed effect, from which estimated marginal means were extracted. To determine whether mortality differed between native and exotic species, we fitted a binomial generalized linear mixed-effects model (GLMM) with a logit link, using counts of dead and surviving trees as the response, and including species as a random intercept. Overall effects of year and origin (native vs exotic), and their interaction, were tested with likelihood ratio tests of fixed effects. To assess origin effects within each year, separate year-specific GLMMs were fitted with species as a random effect, with post-hoc comparisons using compact letter displays where required. To test whether growth differed between native and exotic species, we fitted linear mixed-effects models (LMMs) for each growth metric (e.g., AGR/RGR for height and diameter), with origin, year, and their interaction as fixed effects; species and plant identity were included as random effects to account for repeated measures across taxa. Post-hoc contrasts of estimated marginal means were used to compare origin groups within years, where interactions were significant. We used a one-way ANOVA to test if crown dieback differed between native and exotic species.

Finally, to link demographic outcomes to functional traits, we tested whether species-level survival and crown dieback were associated with measured traits means using trait–outcome regressions. We analysed species’ mean crown dieback against each functional trait using ordinary least squares regression, testing for nonlinearity when ecologically expected. For every trait, we first fitted a linear model (Dieback ∼ trait) before additionally fitting a quadratic model (Dieback ∼ trait + trait²) for water potential metrics (ΔΨ and Ψ_PD_); the best model for each trait was selected using Akaike’s information criterion, or the lowest AIC. We tested for normality using the Shapiro–Wilk test. We extracted slopes, intercepts, coefficients of determination (*r*²), and p-values (p < 0.05). All statistical analyses were conducted in R version 4.4.1 (R Core Team, 2024), using the packages *lme4* for mixed-effects models, *lmer*Test for significance testing, emmeans for estimated marginal means and post hoc comparisons, and ggplot2 for visualisation.

## Results

Climate varied over the study, with the establishment years of 2020-2022 being generally cooler and wetter than the post-establishment years of 2023-2025 (Fig. 2, Table S2). Rainfall was above average in 2020-2022 (935-1498 mm vs 768.5mm, 1993-2025) but 2023 was a dry year with only 532 mm of rain, with intermediate totals in 2024-2025 (829-938 mm). Maximum annual temperatures were about 1-2°C higher in the post-establishment years (25.1-25.5 °C; 2023-2025) than in the establishment years (23.2-24.4 °C; 2020-2022). Differences in maximum monthly temperature were even higher, particularly during summer months – for example, December was ∼4.5 °C hotter in 2023–2025 than in 2020–2022 (31.1-32.6°C vs 26.9-27.8°C, respectively). Together, these patterns indicate several abnormally wet periods during the establishment years before shifting to a hotter climate for the post-establishment years, including periods of low rainfall such as 2023 and the austral 2024-2025 summer (114 mm vs 260.9 mm, 1993-2025).

### Growth, Dieback, and Mortality

There was a large initial decline in survival during the first year after planting, with 27.5% mortality, causing survival to drop to 72.5% (95% CI: 70.2-74.6%). Mortality continued over the next 3.5 years but at a slower annual rate of 7-8% (Table S5). By 2025, survival had decreased to 43.2% (95% CI: 40.7-45.7%; Table 1). Species differed significantly in their survival (χ²₄₇ = 548.5, *p* < 0.001). There were 21 species with relatively high survival (>50%), 17 species with intermediate survival (10–50%) and ten species with critically low survival (<10%) at 4.5 years after planting. Two species, *Tibouchina granulosa* and *Ceratopetalum gummiferum,* had 97% mortality, while another two species, *Pipturus argenteus* and *Xanthostemon chrysanthus*, had complete mortality in 2025 (0% survival; Table S1). Mean first-census mortality was 67.2% in these ten species, compared with 15% in the remaining species, and this difference was highly significant in a quasibinomial GLM (*p* <0.001). Crown dieback differed significantly among species (F_43,2726_ = 8.37, *p* < 0.001) after an extreme heatwave in January 2026, ranging from a mean of 4% in native *Eucalyptus albens* to 95% in exotic *Acer palmatum*. Seven species had high crown dieback (>40%), including *Magnolia grandiflora, Eucalyptus globulus, Pyrus nivalis, Elaeocarpus reticulatus, Quercus dentata, Syzygium luehmannii,* and *Liquidambar styraciflua* (Table S1).

Growth increased most rapidly during the first two years after planting for both height (F_3,2717_ = 427.55, *p* < 0.001) and diameter (F_3, 2737_ = 484.07, *p* < 0.001; Fig. 3). Mean height increased by 63 cm from 2021 to 2022 but only by 31 cm from 2024 to 2025, representing a decrease in growth rate by half. Diameter also increased by 19 mm from 2021 to 2022 but only by 10 mm from 2024 to 2025. Growth from 2021 to 2025 differed significantly among 38 species for both relative and absolute height (RGR: F_37,447 =_9.36, *p* < 0.001; AGR: F₃₇,₄₄₇ = 13.12, *p* < 0.001) and diameter (AGR: F_37,598_ = 9.50, *p* < 0.001; RGR: F_37,598_ = 6.63, *p* < 0.001; Table S1). The species with the fastest mean height and/or diameter growth, which also had the highest absolute height and/or diameter, included *Corymbia eximia, Acacia implexa,* and *E. albens*. In contrast, *M. grandiflora*, *Ficus coronata*, and *Cupaniopsis anacardioides* showed negative height and diameter RGR, consistent with severe stem dieback preventing growth, although basal resprouting was observed in these species (Table S1).

**Figure 3.**
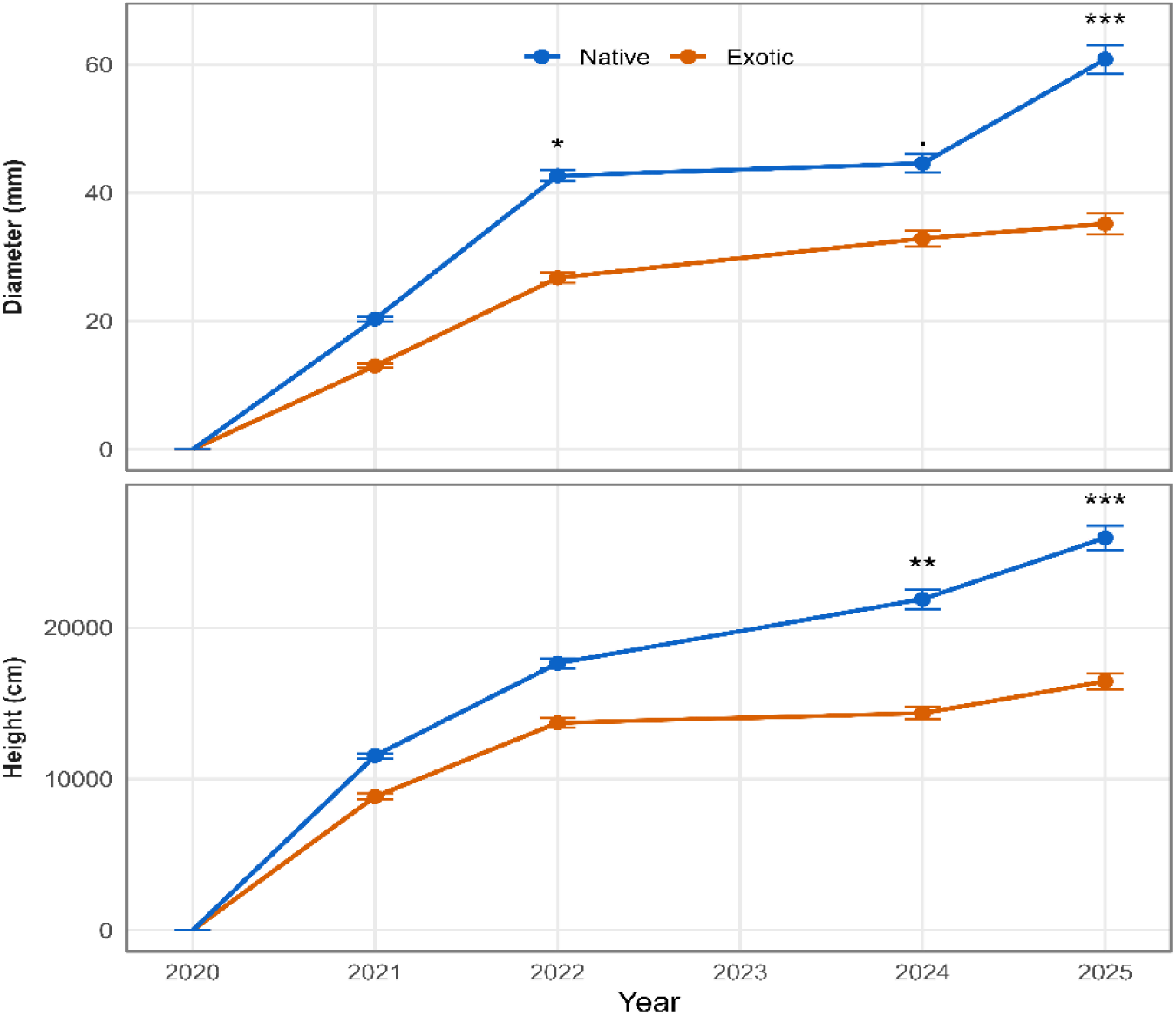
Mean (± SE) changes in tree height and stem diameter for native (red solid line) and exotic (blue dashed line) species across annual inventories (2021, 2022, 2024, 2025). Points show group means pooled across all measured trees within each year, with error bars.

### Differences Between Native and Exotic Species

Exotic species consistently maintained higher survival than native species (χ²₁ = 33.88, *p* < 0.001), but this difference was only significant from 2022 onward (*p* ≤ 0.020; Table 2). Mean survival was 50% for native and 40% for exotic species far more than 10% difference in 2025. Growth differences between native and exotic species changed over time for both height and diameter (Origin × Year: F_3, 2675_ = 26.24, *p* < 0.001; F_3, 2806_ = 9.75, *p* < 0.001, respectively), with native species reaching a larger size than exotic species by the end of the study period (2024–2025; Fig. 3). For height, native species were not different from exotic species in 2021 (Δ = +8 cm, *p* = 0.73) or 2022 (Δ = +28 cm, *p* = 0.22) but were significantly taller in 2024 (Δ = +56 cm, *p* = 0.01) and 2025 (Δ = +69 cm, *p* = 0.002). For diameter, origin differences were non-significant in 2021 (Δ = +6 mm, *p* = 0.28) and marginal in 2024 (Δ = +11 mm, *p* = 0.055) but significant in 2022 (Δ = +13 mm, *p* = 0.019) and 2025 (Δ = +22 mm, *p* < 0.001). From 2021-2025, native species had higher absolute height growth than exotic species (40.0 ± 2.10 vs. 21.4 ± 1.35, respectively), but the difference was marginally non-significant (F_1,39_ = 3.56, *p* = 0.067). Diameter AGR was also only marginally higher for native than exotic species (+4.29 ± 2.42; F_1,37_ = 3.15, *p* = 0.084). Relative growth rates (RGR) did not differ between origin groups for either height (*p* = 0.654) or diameter (*p* = 0.638). Species’ mean crown dieback in January 2026 differed significantly between origin groups, with exotic species having greater mean crown dieback (39%) than native species (22%; F_1,39_ = 4.46, *p* = 0.041).

**Table 2.** Mean survival and mortality of exotic and native species planted at the common garden in western Sydney, Australia, measured at four time points over a 4.5-year period (2021–2025). For each census year, the number of individuals recorded alive, and their corresponding survival percentages (relative to the total 1,536 planted seedlings), are shown. Asterisks denote a statistically significant difference in survival between native and exotic species in that year (* p < 0.05; ** p < 0.01).

| Year | Month | Alive native | Alive exotic | Survival native (%) | Survival exotic (%) |
| --- | --- | --- | --- | --- | --- |
| 2021 | December | 824 | 289 | 71.5 | 75.3 |
| 2022* | December | 719 | 265 | 62.4 | 69 |
| 2024** | April | 541 | 228 | 47 | 59.4 |
| 2025** | May | 468 | 195 | 40.7 | 50.8 |

### Can Plant Functional Traits Predict Survival and Dieback Among Species?

There was substantial interspecific variation in the three hydraulic traits used to characterise water-use behaviour across the the subset of 30 species. Turgor loss point (π_tlp_) ranged from -1.54 MPa in *Eucalyptus rubida* to -2.48 MPa in *Pyrus nivalis*, while wood density varied from 0.37 g cm⁻³ in *M. grandiflora* to 0.80 g cm⁻³ in *C. anacardioides* (Table 1). The peak summer ΔΨ ranged from near zero (-0.09 MPa) in *Syzygium luehmannii* to -2.70 MPa in *P. nivalis* (Table 1). Among all 30 species, the four most isohydric species were all native Australian species, including *S. luehmannii, Acacia implexa, Elaeocarpus reticulatus,* and *Eucalyptus mannifera*. The four most anisohydric species included both native and exotic species: *Callistemon viminalis, Callistemon pallidus, Quercus palustris*, and *P. nivalis* (Table 1).

No plant functional trait was significantly correlated to survival across the subset of 30 species (Table S3). Six plant functional traits were significantly correlated to mean dieback across species (Table S4). Crown dieback was most strongly associated with plant water relations traits, including ΔΨ (*r*² = 0.318, *p* = 0.001; Fig. 4a), Ψ_PD_ (*r*² = 0.297, *p* = 0.002; Fig. 4b), and leaf relative water content (*r*² = 0.256, *p* = 0.004; Fig. 4e) but not π_tlp_ (*r*² = 0.089, *p* = 0.109; Fig. 4c). Relationships between ΨPD and species’ dieback were nonlinear (peaked, quadratic), while leaf RWC was linearly related to dieback across species. Among structural traits, wood density was linearly and negatively correlated to dieback across species (*r*² = 0.207, *p* = 0.012; Fig. 4d), while leaf width (*r*² = 0.203, *p* = 0.012; Fig. 4f) and leaf size (*r*² = 0.163, *p* = 0.027; Table S4) were linearly and positively correlated to dieback across species. There was little evidence for relationships between species dieback and any other measured trait, including survival, growth (height, stem collar diameter), T_leafmax_, stomatal density, midday Ψ_MD_, Φ_PSII_, gs, or leaf thickness (*r*² ≤ 0.073, *p* ≥ 0.149; Table S4).

**Figure 4.**
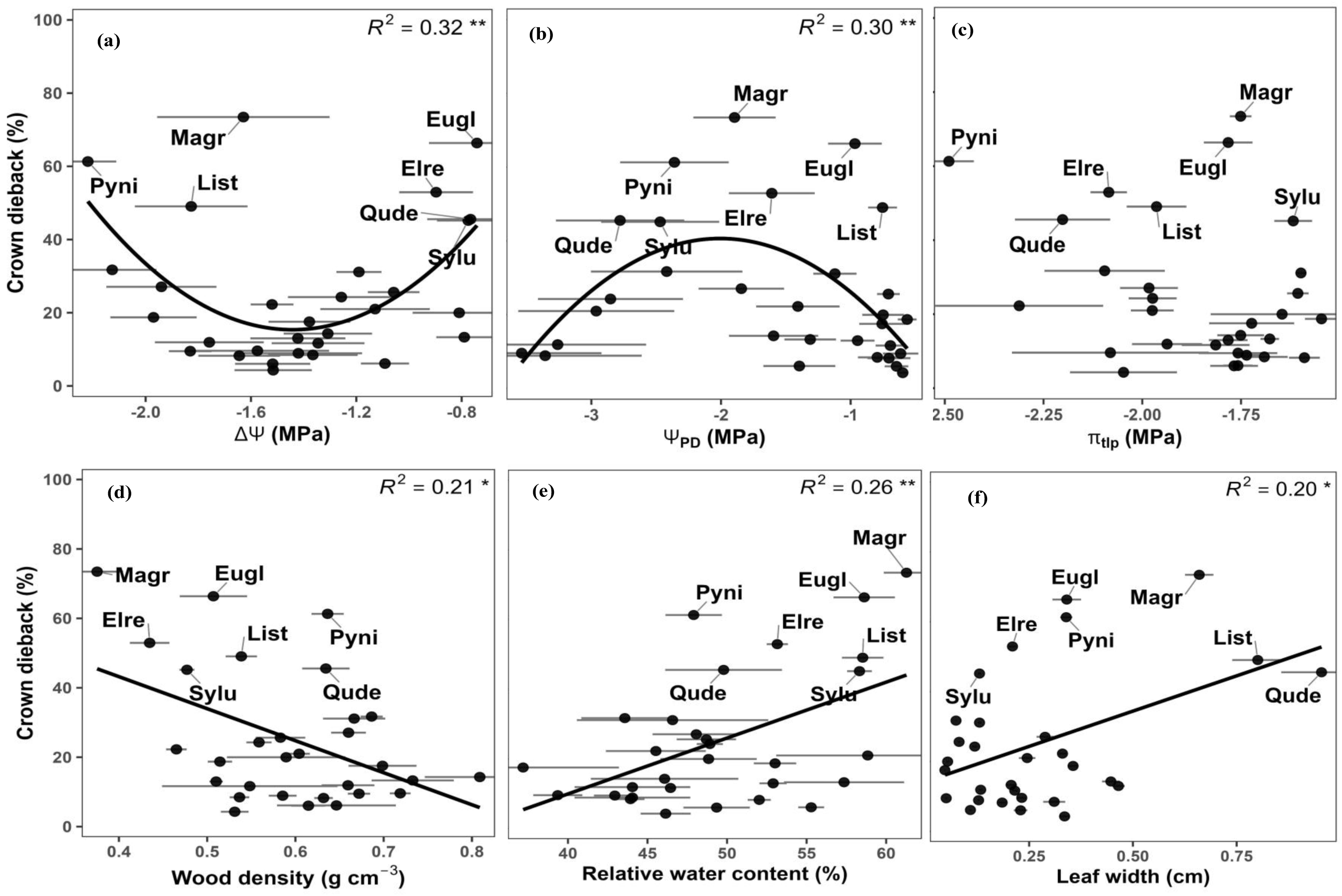
Relationships between mean crown dieback (%) and key leaf functional traits across species in the Hawkesbury Urban Forest common-garden (Richmond, NSW, Australia): Species-level mean crown dieback (%) is plotted against key hydraulic, thermal, and morphological traits: (a) water potential difference ΔΨ (MPa) (b) predawn water potential, (Ψ_PD)_ (MPa); (c) turgor loss point (π_tlp)_ (MPa); (d) wood density (g cm^-3^); (e) relative water content (%); and (f) leaf width (cm). Species codes are as follows: Magr (*Magnolia grandiflora*), Elre (*Elaeocarpus reticulatus*), Eugl (*Eucalyptus globulus*), List (*Liquidambar styraciflua*), Sylu (*Syzygium luehmannii*), Pyni (*Pyrus nivalis*), Qude (*Quercus dentata*). Points represent species means with horizontal and vertical error bars showing ±SE for the trait and dieback, respectively. Solid lines indicate fitted linear relationships; marginal (r^2^) values are shown in each panel, with significance denoted by p < 0.05 (*) and p < 0.01 (**).

## Discussion

Urban forests are increasingly exposed to climate-related stresses, particularly extreme heat and more frequent or intense dry periods, which can interact with local urban constraints such as compacted or low-quality soils (Esperon-Rodriguez et al., 2025a; Kibria et al., 2024). We monitored growth, mortality, and crown dieback of 48 tree or shrub species in a common garden in western Sydney, finding that native species had less crown dieback and higher absolute growth (Fig. 3), but lower survival (Table 2), than exotic species at about five years after planting. Seven Australian native species experienced substantial mortality during the first study year (22-100%), suggesting they may have been poorly suited to local climate conditions, site condition (such as soils), including the periodic flooding that occurred during the establishment years (2020-2022; Kibria et al., 2026). There was a clear native growth advantage by 2025 (Fig. 3), however, after several hotter and drier summers (Table S2). Native species were adapted to local heat extremes, as evidenced by their tendency for higher growth in hotter years (Fig. 3), whereas exotic species accumulated greater dieback than native species after an extreme three-day heatwave >40°C in January 2026. Seven species experienced extensive dieback (mean: >40%) from the compound heat and drought events, including four exotic and three native species (Fig. 4, Table S4). There were no significant correlations between species’ survival in May 2025 and any of the measured plant functional traits (Table S3), however, reinforcing that differential patterns of survival across species were likely not primarily driven by hotter drought stress.

### Native Species Were More Tolerant of Cumulative Thermal Stress

Mean survival declined sharply in the first year after planting by 27.5% but continued gradually decreasing by 7-8% for the next 3.5 years (Table S5). There was a general shift in climate throughout the study, from cooler and wetter conditions in 2021-2022 (Bureau of Meteorology, 2023) to a warmer and drier climate in 2023-2026 (Table S2), exposing trees to contrasting stresses over time. Among the exotic species, *L. styraciflua* had the greatest mean height (252 cm), whereas among the native species, *E. albens* had the greatest mean height (450 cm). Thus, the tallest native species was 198 cm taller on average than the tallest exotic species. The different timing of the observed patterns in mortality, growth, and crown dieback between native and exotic species suggests there may have been different physiological mechanisms involved. Early mortality and establishment failure may have been linked to poor climate suitability. planting site conditions perhaps triggered by the flooding that occurred in 2021 and 2022 (Kibria et al., 2026). Out of the ten species with critically low survival (<10%) in May 2025, seven were native species –four of which were tropical (Kibria et al., 2026). These native tropical species, such as *Xanthostemon chrysanthus, Pipturus argenteus, Tibouchina granulosa* and *Ceratopetalum gummiferum*, were likely intolerant to such conditions (Kibria et al., 2026).

Post-establishment mortality and crown dieback for both native and exotic species was likely more closely related to periodic drought stress and associated cumulative thermal stress in 2023-2026. Consistent with this, several heat- and drought-tolerance traits were correlated with dieback across species, including leaf size, leaf width, predawn leaf water potential, daily water potential difference (ΔΨ), wood density, and relative water content (Fig. 4; Table S4). Notably, species’ crown dieback was not correlated to their survival (Tables S3, S4), suggesting that the factors causing mortality were not entirely the same as those causing crown dieback in January 2026. The correlations between crown dieback and plant functional traits provide evidence that heat and drought stress is the cause of the ongoing gradual 6-8% annual mortality throughout the post-establishment years. Three exotic species, *Q. dentata, L. styraciflua* and *M. grandiflora*, had the largest leaves, along with extensive crown dieback (Fig. 4f). Large leaves have thick boundary layers, which reduce convective heat loss and increase leaf temperature under strong radiation and stomatal closure (Leigh et al., 2012, 2017; Vogel, 2009). Crown dieback was also high in species with low wood density, including one exotic (*M. grandiflora*) and three native species (*E. reticulatus*, *S. luehmannii*, *E. globulus*). Low wood density is often associated with lower drought resistance and greater susceptibility to xylem cavitation or cell wall collapse (Hacke et al., 2001; Greenwood et al., 2017; Santiago et al., 2004; Kraft et al., 2010; Nascimento et al., 2005). This association is not universal, as wood density is not always tightly linked to embolism resistance across species, particularly for species with hydraulic vulnerability that depends more directly on pit membrane structure (Petruzzellis et al., 2021; Trueba et al., 2017; Delzon et al., 2010). A previous study on urban street trees growing in western Sydney also found that species with low wood density were vulnerable to dieback and mortality during an extreme summer (Marchin et al., 2022a). Taken together with the wide variation among species within both origin groups, our results suggest that post-establishment performance at our study site was shaped more by cumulative thermal stress than by climate of origin alone (Franklin et al., 1987; Czaja et al., 2020; Hauer, Koeser, et al., 2020b; Hilbert et al., 2019). Plant functional traits were more informative than broad native-exotic origin contrasts for predicting urban tree stress tolerance and performance (Kendal et al., 2014; Sjöman et al., 2016; Marchin et al., 2022a).

### Intermediate, rather than Isohydric or Anisohydric Species, Avoided Dieback

Crown dieback after the extreme heatwave was most strongly correlated to plant water relations traits, including predawn water potential (Ψ_PD_; Fig. 4b) and ΔΨ (Fig. 4a), each explaining about 30% of the variation in crown dieback among species. The Ψ_PD_ represents plant available water, while ΔΨ is the daily decline in leaf water potential and indicates daily water use along with species’ degree of isohydricity (Martínez-Vilalta et al., 2014). We found that species with extensive crown dieback all had either low or high ΔΨ (Fig. 4a), suggesting both isohydric and anisohydric species were vulnerable to compound heat and drought stress at our site. For example, two of most isohydric (*S. luehmannii*, *E. reticulatus*) and two of the most anisohydric (*P. nivalis*, *Q. dentata*) species were vulnerable with high crown dieback (Table S1, Fig. 4). Isohydric species had low wood density and/or high TLP, along with small ΔΨ, while anisohydric species had high wood density, low TLP, and/or large ΔΨ (Table 1, Fig. 4). High crown dieback of both isohydric and anisohydric species may have been caused by catastrophic xylem embolism and hydraulic failure. Isohydric species have low drought tolerance, and the Ψ_PD_ of at least one isohydric species (*S. luehmannii*) was below their TLP in the austral 2024-2025 summer (Fig. 4). Despite having more resistant xylem to cavitation, anisohydric species tend to maintain smaller hydraulic safety margins and therefore can also be vulnerable during drought (Skelton et al., 2015; Anderegg et al., 2015, Hoffmann et al., 2011). Here, the Ψ_PD_ of anisohydric species *Q. dentata* was also below their TLP (Fig. 4). The cause for crown dieback in other vulnerable species is less certain but is likely related to cumulative thermal stress, at least for exotic species with large leaves such as *L. styraciflua* and *M. grandiflora* (Fig. 4f).

Interestingly, vulnerability to compound heat and drought stress at our study site did not fall neatly into a single hydraulic strategy group. Intermediate ‘mesohydric’ species, defined as those exhibiting intermediate hydraulic regulation between the isohydric to anisohydric continuum, such as *F. griffithii, Q.ilex, F. oxycarpa, U. parvifolia*, and *F. australis* avoided extensive dieback (6.1-21.0%; Table S1). This agrees with Gu et al. (2015), who argued that mortality risk may vary nonlinearly across the isohydric-anisohydric continuum, although other studies have found that either isohydric or anisohydric behaviour is favoured depending on site and climate context (Feng et al., 2019; Hoffmann et al., 2011). For example, anisohydric species have been vulnerable in a temperate forest under severe drought (Hoffmann et al., 2011), whereas isohydric species were previously found to be vulnerable in western Sydney’s urban forest during a heatwave (Marchin et al., 2022a). Together, these results indicate that crown dieback was a multi-causal process, but one strongly associated with leaf energy balance, hydraulic stress, and structural resistance during the hotter and drier post-establishment years.

### Species Selection for Urban Forests

Two species, *C. eximia* and *A. implexa*, had among the highest values for both height RGR and AGR, indicating rapid growth both relative to their initial size and in absolute terms. Two *Banksia* species, *B. ericifolia* and *B. integrifolia*, also had high RGR, whereas two of the *Eucalyptus* species, *E. rubida* and *E. albens*, had high AGR. Rapid growth did not consistently correspond with high survival, however, suggesting that growth rate alone was not sufficient to confer long-term resilience under multiple stresses (Poorter et al., 2012; Wright et al., 2010). High absolute growth may enhance canopy development and associated ecosystem services, but it may also increase water demand and potentially heighten vulnerability to heat and drought stress (McDowell et al., 2008; Esperon-Rodriguez et al., 2022). These six native species are recommended for planting in Sydney, due to their high growth and survival, combined with low crown dieback (mean: <20%) after the extreme heatwave, suggesting a moderate to high heat tolerance.

Species with limited or negative growth experienced chronic stress at our study site, including *S. luehmannii*, *M. grandiflora*, and *C. anacardioides*, which all had evidence of repeated aboveground stem dieback (mean: 14 -73%) requiring basal resprouting. Similarly, poor diameter growth in *S*. *glomulifera*, *S. australe*, *F. australis*, and *F. coronata* suggests intolerance to heat and drought stress, so these species are not recommended for planting in Sydney unless supplemental irrigation is available. *F. coronata* was the only species with negative values across both AGR and RGR, indicating particularly poor performance. Field observations during winter suggested that frost sensitivity may have contributed to this poor growth response, although frost tolerance was not directly quantified in this study. These results show that growth trends provide important information, alongside crown dieback and survival, for assessing urban tree performance and aiding in selection of species that will thrive under future climates (Franklin et al., 1987; Czaja et al., 2020; Hilbert et al., 2019; Hauer et al., 2020a).

## Conclusion

Our results suggest that urban tree resilience in western Sydney is better explained by species-specific functional traits than by the broader native–exotic classification. Although exotic species achieved higher survival, their growth was generally lower and dieback during an extreme heatwave was higher than that of native species. For example, the tallest exotic species, *L. styraciflua*, was on average ∼2 m shorter than the tallest native species, *E. albens*, suggesting that native species may provide greater ecosystem services than exotic species under western Sydney conditions. This reinforces the preference for native species in some cities but shifting from origin- or climate-based species selection to trait-informed species selection can better match species physiological tolerances and performance with planting contexts. Here, we found ‘mesohydric’ species, here defined as those exhibiting intermediate hydraulic regulation between the isohydric to anisohydric continuum, were more resilient to cumulative thermal stress. Our study also highlights crown dieback as a useful early indicator of stress, with clear value for monitoring and intervention before mortality occurs. In the context of intensifying heatwaves and drought, combining trait-based assessment with long-term field evaluation will be critical for improving species selection and supporting more climate-resilient urban forests.

## Supporting information

Table S1. Summary of tree and shrub species planted in the Hawkesbury Urban Forest common-garden trial (Richmond, NSW, Australia), with family, origin

## Conflict of interest

The authors declare that there is no conflict of interest.

## Acknowledgments

We sincerely thank Pieter A. Arnold and John Gray for their valuable support and assistance throughout this research. This research was supported by the Commonwealth through an Australian Government Research Training Program Scholarship [DOI: https://doi.org/10.82133/C42F-K220] and by the Australian Government through the Australian Research Council Discovery Early Career Researcher Award awarded to Dr R. Marchin (Project DE200100649).

## Notes

### Competing Interest Statement

The authors have declared no competing interest.

## References

Anderegg, W. R. L., Flint, A., Huang, C. Y., Flint, L., Berry, J. A., Davis, F. W., Sperry, J. S., & Field, C. B. (2015). Tree mortality predicted from drought-induced vascular damage. Nature Geoscience, 8(5), 367–371. 10.1038/ngeo2400

Australian Bureau of Meteorology. (2025). Richmond monthly weather data. Commonwealth of Australia. http://www.bom.gov.au/climate/data/ (Accessed 14 October 2024)

Baró, F., Chaparro, L., Gómez-Baggethun, E., Langemeyer, J., Nowak, D. J., & Terradas, J. (2014). Contribution of ecosystem services to air quality and climate change mitigation policies: The case of urban forests in Barcelona, Spain. Ambio, 43(4), 466–479. 10.1007/s13280-014-0507-x

Barros, F. D. V., Bittencourt, P. L., Eller, C. B., Signori-Müller, C., Meireles, L. D., & Oliveira, R. S. (2022). Phytogeographical origin determines Tropical Montane Cloud Forest hydraulic trait composition. Functional Ecology, 36, 607–621. 10.1111/1365-2435.14008

Bartlett, M. K., Klein, T., Jansen, S., Choat, B., & Sack, L. (2016). The correlations and sequence of plant stomatal, hydraulic, and wilting responses to drought. Proceedings of the National Academy of Sciences, 113(46), 13098–13103. 10.1073/pnas.1604088113

Bartlett, M. K., Scoffoni, C., Ardy, R., Zhang, Y., Sun, S., Cao, K., & Sack, L. (2012). Rapid determination of comparative drought tolerance traits: Using an osmometer to predict turgor loss point. Methods in Ecology and Evolution, 3(5), 880–888. 10.1111/j.2041-210X.2012.00230.x

Bottalico, F., Pesola, L., Vizzarri, M., Antonello, L., Barbati, A., Chirici, G., Corona, P., Cullotta, S., Garfì, V., Giannico, V., Lafortezza, R., Lombardi, F., Marchetti, M., Nocentini, S., Puletti, N., Santopuoli, G., Sarao, A., & Ferrini, F. (2017). A spatially-explicit method to assess the dry deposition of air pollutants by urban forests in Mediterranean cities. Urban Forestry & Urban Greening, 26, 163–171. 10.1016/j.ufug.2017.06.013

Brandt, L., Lewis, A. D., Fahey, R., Scott, L., Darling, L., & Swanston, C. (2016). A framework for adapting urban forests to climate change. Environmental Science & Policy, 66, 393–402. 10.1016/j.envsci.2016.06.005

Bureau of Meteorology. (2023, February 7). Greater Sydney in 2022: Wettest year on record for many areas, cooler than average. https://www.bom.gov.au/climate/current/annual/nsw/archive/2022.sydney.shtml

Burley, H., Beaumont, L. J., Ossola, A., Baumgartner, J. B., Gallagher, R., Laffan, S., … & Leishman, M. R. (2019). Substantial declines in urban tree habitat predicted under climate change. Science of the Total Environment, 685, 451–462. 10.1016/j.scitotenv.2019.05.287

Cabral, I., Keim, J., Engelmann, R., Kraemer, R., Siebert, J., Bonn, A., 2017. Ecosystem services of allotment and community gardens: a Leipzig, Germany case study. Urban For. Urban Green. 23, 44–53. 10.1016/j.ufug.2017.02.008

Chen, M., Dang, A., Li, X., Huang, J., & Weng, Y. (2024). Identifying climate-ready plant for urban environment: Integrating machine learning with traditional plant selection tools. Urban Forestry & Urban Greening, 102, 128559. 10.1016/j.ufug.2024.128559

Cook, A. M., Sumner, E. E., Leigh, A., O’Sullivan, O. S., & Curtis, E. M. (2024). Beyond a single temperature threshold: Applying a cumulative thermal stress framework to plant heat tolerance. Ecology Letters, 27(4), e14416. 10.1111/ele.14416

Csete, A.K., ‘ Gulý as, A., ‘ 2021. Green infrastructure-based hydrological modelling, a comparison between different urban districts, through the case of Szeged, Hungary. Hung. Geogr. Bull. 70 (4), 353–368. 10.15201/HUNGEOBULL.70.4.5.

Cuce, P. M., Cuce, E., & Santamouris, M. (2025). Towards Sustainable and Climate-Resilient Cities: Mitigating Urban Heat Islands Through Green Infrastructure. Sustainability, 17(3), 1303. 10.3390/su17031303

Curtis, E. M., Knight, C. A., Petrou, K., & Leigh, A. (2014). A comparative analysis of photosynthetic recovery from thermal stress: A desert plant case study. Oecologia, 175(4), 1051–1061. 10.1007/s00442-014-2988-5

Czaja, M., Kołton, A., & Muras, P. (2020). The complex issue of urban trees—Stress factor accumulation and ecological service possibilities. Forests, 11(9), 932. 10.3390/f11090932

Deb, J. C., Phinn, S., Butt, N., & McAlpine, C. A. (2018). Climate change impacts on tropical forests: Identifying risks for tropical Asia. Journal of Tropical Forest Science, 30(2), 182–194. 10.26525/jtfs2018.30.2.182194

Drago, M., Petrescu, A., Merciu, G.L., 2017. Analysis of vegetation from satelite images correlated to the bird species presence and the state of health of the ecosystems of Bucharest during the period from 1991 to 2006. Geogr. Pannonica 21 (1), 9–25. 10.5937/GEOPAN1701009D

Drake, J. E., Tjoelker, M. G., Vårhammar, A., Medlyn, B. E., Reich, P. B., Leigh, A., Pfautsch, S., Blackman, C. J., López, R., Aspinwall, M. J., Crous, K. Y., Duursma, R. A., Kumarathunge, D., De Kauwe, M. G., Jiang, M., Nicotra, A. B., Tissue, D. T., Choat, B., Atkin, O. K., & Barton, C. V. M. (2018). Trees tolerate an extreme heatwave via sustained transpirational cooling and increased leaf thermal tolerance. Global Change Biology, 24(6), 2390–2402. 10.1111/gcb.14037

Esperon-Rodriguez, M., Gallagher, R. V., Souverijns, N., Lejeune, Q., Schleussner, C. F., & Tjoelker, M. G. (2024a). Mapping the climate risk to urban forests at city scale. Landscape and Urban Planning, 248, 105090. 10.1016/j.landurbplan.2024.105090

Esperon-Rodriguez, M., Gallagher, R., Calfapietra, C., Cariñanos, P., Dobbs, C., Eleuterio, A. A., … & Tjoelker, M. G. (2025a). Barriers and opportunities for resilient and sustainable urban forests. Nature Cities, 2(4), 290–298. 10.1038/s44284-025-00100-y

Esperon-Rodriguez, M., Power, S. A., Tjoelker, M. G., Beaumont, L. J., Burley, H., Caballero-Rodriguez, D., & Rymer, P. D. (2019). Assessing the vulnerability of Australia’s urban forests to climate extremes. Plants, People, Planet, 1(4), 387–397. 10.1002/ppp3.10052

Esperon-Rodriguez, M., Power, S. A., Tjoelker, M. G., Marchin, R. M., & Rymer, P. D. (2021). Contrasting heat tolerance of urban trees to extreme temperatures during heatwaves. Urban Forestry & Urban Greening, 66, 127387. 10.1016/j.ufug.2021.127387

Esperon-Rodriguez, M., Power, S. A., Tjoelker, M. G., Sharmin, M., & Rymer, P. D. (2025b). A nation-wide urban trial network of tree and shrub climate resilience. Urban Forestry & Urban Greening, 105, 128720. 10.1016/j.ufug.2024.128720

Esperon-Rodriguez, M., Rymer, P. D., Power, S. A., Barton, D. N., Cariñanos, P., Dobbs, C., … & Tjoelker, M. G. (2022). Assessing climate risk to support urban forests in a changing climate. Plants, People, Planet, 4(3), 201–213. 10.1002/ppp3.1235

Esperon-Rodriguez, M., Tjoelker, M. G., Lenoir, J., et al. (2024b). Climate-resilient urban forests require species-specific evidence-based planning. Nature Climate Change, 14, 245–252. 10.1038/s41558-024-01936-x

Farrell, C., Szota, C., Arndt, S. K., & Livesley, S. J. (2022). Can we integrate ecological approaches to improve plant selection for green infrastructure? Urban Forestry & Urban Greening, 73, 127617. 10.1016/j.ufug.2022.127617

Feng, X., Ackerly, D. D., Dawson, T. E., Manzoni, S., McLaughlin, B., Skelton, R. P., Vico, G., Weitz, A. P., & Thompson, S. E. (2019). Beyond isohydricity: The role of environmental variability in determining plant drought responses. Plant, Cell & Environment, 42(4), 1104–1111. 10.1111/pce.13486

Franceschi, E., Moser-Reischl, A., Honold, M., Rahman, M. A., Pretzsch, H., Pauleit, S., & Rötzer, T. (2023). Urban environment, drought events and climate change strongly affect the growth of common urban tree species in a temperate city. Urban Forestry & Urban Greening, 88, 128083.

Franceschi, M., Beaumont, L. J., & Esperon-Rodriguez, M. (2023). Widespread leaf scorching and premature leaf drop in urban trees during a record-breaking heatwave. Urban Forestry & Urban Greening, 86, 128031. 10.1016/j.ufug.2023.128031

Franklin, J. F., Shugart, H. H., & Harmon, M. E. (1987). Tree death as an ecological process. Bioscience, 37(8), 550–556. 10.2307/1310665

Friess, D.A., Richards, D.R., Phang, V.X.H., 2016. Mangrove forests store high densities of carbon across the tropical urban landscape of Singapore. Urban Ecosyst. 19 (2), 795–810. 10.1007/s11252-015-0511-3

Greenwood, S., Ruiz-Benito, P., Martínez-Vilalta, J., Enquist, B. J., Jump, A. S., Scott, S. L., … & Zavala, M. A. (2017). Tree mortality across biomes is promoted by drought intensity, lower wood density and higher specific leaf area. Ecology Letters, 20(4), 539–553. 10.1111/ele.12748

Gu, L., Pallardy, S. G., Hosman, K. P., & Sun, Y. (2015). Predictors and mechanisms of the drought-influenced mortality of tree species along the isohydric to anisohydic continuum in a decade-long study of a central US temperate forest. Biogeosciences Discussions, 12(2).

Hacke, U.G., Sperry, J.S., Pockman, W.T., Davis, S.D., McCulloh, K.A., 2001. Trends in wood density and structure are linked to prevention of xylem implosion by negative pressure. Oecologia 126, 457–461.

Hauer, R., Hanou, I., & Sivyer, D. (2020a). Planning for active management of future invasive pests affecting urban forests: The ecological and economic effects of varying Dutch elm disease management practices for street trees in Milwaukee, WI USA. Urban Ecosystem, 23(5), 1005–1022. 10.1007/s11252-020-00976-6

Hauer, R., Koeser, A., Parbs, S., Kringer, J., Krouse, R., Ottman, K., Miller, R., Sivyer, D., Timilsina, N., & Werner, L. (2020b). Effects of a tree preservation program on tree survival, condition, and growth in Milwaukee, WI, USA. Landscape & Urban Planning, 193, 103670. 10.1016/j.landurbplan.2019.103670

Hilbert, D., Roman, L., Koeser, A. K., Vogt, J., & Van Doorn, N. S. (2019). Urban Tree Mortality: A Literature Review. Arboriculture & Urban Forestry, 45(5), 167–200.

Hoffmann, W. A., Marchin, R. M., Abit, P., & Lau, O. L. (2011). Hydraulic failure and tree dieback are associated with high wood density in a temperate forest under extreme drought. Global Change Biology, 17(8), 2731–2742.

Huang, S., Knight, C. A., Hoover, B. K., & Ritter, M. (2020). Leaf functional traits as predictors of drought tolerance in urban trees. Urban Forestry & Urban Greening, 48, 126577. 10.1016/j.ufug.2019.126577

Hüve, K., Bichele, I., Rasulov, B., & Niinemets, Ü. (2011). When it is too hot for photosynthesis: Heat-induced instability of photosynthesis in relation to respiratory burst, cell permeability changes and H2O2 formation. Plant, Cell & Environment, 34(1), 113–126. 10.1111/j.1365-3040.2010.02229.x French, K., Jansens, I. B., Ashcroft, M. B., Ecroyd, H., & Robinson, S. A. (2019). High tolerance of repeated heatwaves in Australian native plants. Austral Ecology, 44(4), 597–608. 10.1111/aec.12707

Keeler, B. L., Hamel, P., McPhearson, T., Hamann, M. H., Donahue, M. L., Meza Prado, K. A., & Wood, S. A. (2019). Social-ecological and technological factors moderate the value of urban nature. Nature Sustainability, 2(1), 29–38.

Kendal, D., Williams, K. J. H., & Williams, N. S. G. (2014). Plant traits link people’s plant preferences to the composition of their gardens. Landscape and Urban Planning, 124, 34–42. 10.1016/j.landurbplan.2014.01.002

Kibria, M. G., Tjoelker, M. G., Esperón-Rodríguez, M., Marchin, R. M., Arndt, S. K., Livesley, S. J., & Rymer, P. D. (2026). Functional traits are better than climate niche approach at predicting the vital growth and mortality rates of urban tree species. Urban Forestry & Urban Greening, 120, 129450. 10.1016/j.ufug.2026.129450

Kibria, M. G., Tjoelker, M. G., Marchin, R. M., Arndt, S. K., & Rymer, P. D. (2024). Can species climate niche predict canopy growth, functional traits and phenotypic plasticity in urban trees? Urban Forestry & Urban Greening, 98, 128417. 10.1016/j.ufug.2024.128417

Kirilenko, A. P., & Sedjo, R. A. (2007). Climate change impacts on forestry. Proceedings of the National Academy of Sciences of the United States of America, 104(50), 19697–19702. 10.1073/pnas.0701424104

Klein, T. (2014), The variability of stomatal sensitivity to leaf water potential across tree species indicates a continuum between isohydric and anisohydric behaviours. Funct Ecol, 28: 1313–1320. 10.1111/1365-2435.12289

Kraft, N. J., Metz, M. R., Condit, R. S., & Chave, J. (2010). The relationship between wood density and mortality in a global tropical forest data set. New Phytologist, 188(4), 1124–1136. 10.1111/j.1469-8137.2010.03444.x

Leigh, A., Sevanto, S., Ball, M. C., Close, J. D., Ellsworth, D. S., Knight, C. A., & Vogel, S. (2012). Do thick leaves avoid thermal damage in critically low wind speeds? New Phytologist, 194(2), 477–487. 10.1111/j.1469-8137.2012.04054.x

Lindner, M., Maroschek, M., Netherer, S., Kremer, A., Barbati, A., Garcia-Gonzalo, J., Seidl, R., Delzon, S., Corona, P., Kolström, M., Lexer, M. J., & Marchetti, M. (2010). Climate change impacts, adaptive capacity, and vulnerability of European forest ecosystems. Forest Ecology and Management, 259(4), 698–709. 10.1016/j.foreco.2009.09.023

Liu, J., Zhang, L., Zhang, Q., Zhang, G., Teng, J., 2021. Predicting the surface urban heat island intensity of future urban green space development using a multi-scenario simulation (Article). Sustain. Cities Soc. 66, 102698. 10.1016/j.scs.2020.102

Liu, Z., He, C., Zhou, Y. & Wu, J. How much of the world’s land has been urbanized, really? A hierarchical framework for avoiding confusion. Landsc. Ecol. 29, 763–771 (2014).

Manoli, G., Fatichi, S., Schläpfer, M., Yu, K., Crowther, T. W., Meili, N., Burlando, P., Katul, G. G., & Bou-Zeid, E. (2019). Magnitude of urban heat islands largely explained by climate and population. Nature, 573(7772), 55–60. 10.1038/s41586-019-1512-9

Marchin, R. M., Backes, D., Ossola, A., Leishman, M. R., Tjoelker, M. G., & Ellsworth, D. S. (2022b). Extreme heat increases stomatal conductance and drought-induced mortality risk in vulnerable plant species. Global Change Biology, 28(3), 1133–1146. 10.1111/gcb.15967

Marchin, R. M., Esperon-Rodriguez, M., Tjoelker, M. G., & Ellsworth, D. S. (2022a). Crown dieback and mortality of urban trees linked to heatwaves during extreme drought. Science of The Total Environment, 850, 157915. 10.1016/j.scitotenv.2022.157915

Maréchaux, I., Bartlett, M. K., Sack, L., Baraloto, C., Engel, J., Joetzjer, E., & Chave, J. (2015). Drought tolerance as predicted by leaf water potential at turgor loss point varies strongly across species within an Amazonian forest. Functional Ecology, 29(10), 1268–1277. 10.1111/1365-2435.12452

Martin, N.A., Chappelka, A.H., Loewenstein, E.F., Keever, G.J., 2012. Comparison of carbon storage, carbon sequestration, and air pollution removal by protected and maintained urban forests in Alabama, USA. Int. J. Biodivers. Sci. Ecosyst. Serv. Manag. 8 (3), 265–272. 10.1080/21513732.2012.712550.

Martínez-Vilalta, J., Poyatos, R., Aguadé, D., Retana, J., & Mencuccini, M. (2014). A new look at water transport regulation in plants. New Phytologist, 204(1), 105–115. 10.1111/nph.12912

Mccaig, T. N., and Romagosa, I. (1989). Measurement and use of excised-leaf water status in wheat. Crop Sci. 29, 1140–1145. doi: 10.2135/cropsci1989.0011183X002900050008x

McDowell, N., Pockman, W. T., Allen, C. D., et al. (2008). Mechanisms of plant survival and mortality during drought: Why do some plants survive while others succumb? New Phytologist, 178, 719–739.

MEA. 2005b. Ecosystems and Their Services (Ecosystems and Human Well-being: A Framework for Assessment, Issue. https://www.millenniumassessment.org/documents/document.300.aspx.pdf

Michaletz, S. T., Weiser, M. D., McDowell, N. G., Zhou, J., Kaspari, M., Helliker, B. R., & Enquist, B. J. (2016). The energetic and carbon economic origins of leaf thermoregulation. Nature plants, 2(9), 16129.

Moody, R., Geron, N., Healy, M., Rogan, J., Martin, D., 2021. Modeling the spatial distribution of the current and future ecosystem services of urban tree planting in G. Sharma et al. Urban Forestry & Urban Greening 104 (2025) 128659 11 Chicopee and Fall River, Massachusetts. Urban For Urban Green. 66, 127403. 10.1016/j.ufug.2021.127403.

Nascimento, H. E., Laurance, W. F., Condit, R., Laurance, S. G., D’Angelo, S., & Andrade, A. C. (2005). Demographic and life-history correlates for Amazonian trees. Journal of Vegetation Science, 16(6), 625–634.

Neuner, G., & Buchner, O. (2023). The dose makes the poison: the longer the heat lasts, the lower the temperature for functional impairment and damage. Environmental and Experimental Botany, 212, 105395.

Nowak, D. J., & Crane, D. E. (2002). Carbon storage and sequestration by urban trees in the USA. Environmental Pollution, 116(3), 381–389.

O’Brien, M. J., Engelbrecht, B. M., Joswig, J., Pereyra, G., Schuldt, B., Jansen, S., … & Leuzinger, S. (2017). A synthesis of tree functional traits related to drought-induced mortality in field studies. Global Change Biology, 23(9), 3731–3743. 10.1111/gcb.13651

Oke, T. R. (1982). The energetic basis of the urban heat island. Quarterly journal of the royal meteorological society, 108(455), 1–24.

Petri, A. C., Koeser, A. K., Lovell, S. T., & Ingram, D. (2016). How green are trees?—Using life cycle assessment methods to assess net environmental benefits. Journal of Environmental Horticulture, 34(4), 101–110.

Petruzzellis, F., Tordoni, E., Di Bonaventura, A., Tomasella, M., Natale, S., Panepinto, F., … & Nardini, A. (2022). Turgor loss point and vulnerability to xylem embolism predict species-specific risk of drought-induced decline of urban trees. Plant Biology, 24(7), 1198–1207. 10.1111/plb.13454

Poorter, H., Niklas, K. J., Reich, P. B., Oleksyn, J., Poot, P., & Mommer, L. (2012). Biomass allocation to leaves, stems and roots: Meta-analyses of interspecific variation and environmental control. New Phytologist, 193(1), 30–50. 10.1111/j.1469-8137.2011.03952.x

Potgieter, L.J., Gaertner, M., O’Farrell, P.J., Richardson, D.M., 2019. A fine-scale assessment of the ecosystem service-disservice dichotomy in the context of urban ecosystems affected by alien plant invasions. For. Ecosyst. 6 (1), 46. 10.1186/s40663-019-0200-4

Pregitzer, C. C., Hanna, C., Charlop-Powers, S., & Bradford, M. A. (2022). Estimating carbon storage in urban forests of New York City. Urban Ecosystems, 25(2), 617–631. 10.1007/s11252-021-01188-7

Rahman, A. A.-N. S., Rahman, M., Shimanto, M. H., Kibria, M. G., & Islam, M. (2022). Stomatal size and density trade-off varies with leaf phenology and species shade tolerance in a South Asian moist tropical forest. Functional Plant Biology, 49(4), 307–318. 10.1071/FP21255

Rezende, E. L., Castañeda, L. E., & Santos, M. (2014). Tolerance landscapes in thermal ecology. Functional Ecology, 28(4), 799–809. 10.1111/1365-2435.12268

Santamouris, M. (2020). Recent progress on urban overheating and heat island research. Integrated assessment of the energy, environmental, vulnerability and health impact. Synergies with the global climate change. Energy and Buildings, 207, 109482. 10.1016/j.enbuild.2019.109482

Santiago, L. S., Goldstein, G., Meinzer, F. C., Fisher, J. B., Machado, K., Woodruff, D., & Jones, T. (2004). Leaf photosynthetic traits scale with hydraulic conductivity and wood density in Panamanian forest canopy trees. Oecologia, 140(4), 543–550. 10.1007/s00442-004-1624-1

Sanusi, R., & Livesley, S. J. (2020). London Plane trees (Platanus × acerifolia) before, during and after a heatwave: Losing leaves means less cooling benefit. Urban Forestry & Urban Greening, 54, 126746.

Sastry, A., & Barua, D. (2018). Leaf thermotolerance in dry tropical forest tree species: Relationships with leaf traits and effects of drought. AoB Plants, 10(1), plx070. 10.1093/aobpla/plx070

Schneider, C. A., Rasband, W. S., & Eliceiri, K. W. (2012). NIH Image to ImageJ: 25 years of image analysis. Nature Methods, 9(7), 671–675. 10.1038/nmeth.2089

Sjöman, H., Hirons, A. D., & Bassuk, N. L. (2016). Improving confidence in tree species selection for challenging urban sites: A role for leaf turgor loss. Urban Forestry & Urban Greening, 18, 143–151. 10.1016/j.ufug.2016.06.007

Sjöman, H., Watkins, H., Kelly, L. J., Hirons, A., Kainulainen, K., Martin, K. W. E., & Antonelli, A. (2024). Resilient trees for urban environments: The importance of intraspecific variation. Plants, People, Planet, 6(6), 1180–1189. 10.1002/ppp3.10518

Skelton, R. P., West, A. G., & Dawson, T. E. (2015). Predicting plant vulnerability to drought in biodiverse regions using functional traits. Proceedings of the National Academy of Sciences, 112(18), 5744–5749. 10.1073/pnas.1503376112

Slik, J. W. F., Breman, F. C., Bernard, C., van Beek, M., Galley, K. A. O., Keßler, P. J. A., Kartawinata, K., & Sheil, D. (2007). Environmental correlates of tree biomass, basal area, wood specific gravity and stem density gradients in Borneo’s tropical forests. Global Ecology and Biogeography, 16(1), 50–60. 10.1111/j.1466-8238.2006.00265.x

Smith, I. A., Dearborn, V. K., & Hutyra, L. R. (2019). Live fast, die young: Accelerated growth, mortality, and turnover in street trees. PLOS ONE, 14(5), e0215846.

Steenberg, J. W. N., Millward, A. A., Nowak, D. J., & Robinson, P. J. (2017). A conceptual framework of urban forest ecosystem vulnerability. Environmental Reviews, 25(1), 115–126. 10.1139/er-2016-0022

Tabassum, S., Manea, A., Ossola, A., Thomy, B., Blackham, D., & Leishman, M. R. (2021). The angriest summer on record: Assessing canopy damage and economic costs of an extreme climatic event. Urban Forestry & Urban Greening, 63, 127221. 10.1016/j.ufug.2021.127221

Tardieu, F., & Simonneau, T. (1998). Variability among species of stomatal control under fluctuating soil water status and evaporative demand: modelling isohydric and anisohydric behaviours. Journal of experimental botany, 419–432.

Trueba, S., Pouteau, R., Lens, F., Feild, T. S., Isnard, S., Olson, M. E., & Delzon, S. (2017). Vulnerability to xylem embolism as a major correlate of the environmental distribution of rain forest species on a tropical island. Plant, Cell & Environment, 40(2), 277–289.

United Nations, Department of Economic and Social Affairs, Population Division. (2019). World urbanization prospects: The 2018 revision. United Nations.

Van der Veken, S., Hermy, M., Vellend, M., Knapen, A., & Verheyen, K. (2008). Garden plants get a head start on climate change. Frontiers in Ecology and the Environment, 6(4), 212–216. 10.1890/070063

Vogel, S. (2009). Leaves in the lowest and highest winds: temperature, force and shape. New Phytologist, 183(1), 13–26.

Wang, C., He, J., Zhao, T. H., Cao, Y., Wang, G., Sun, B., … & Li, M. H. (2019). The smaller the leaf is, the faster the leaf water loses in a temperate forest. Frontiers in plant science, 10, 58.

Wang, C., Wang, Z. H., & Yang, J. (2018). Cooling effect of urban trees on the built environment of contiguous United States. Earth’s Future, 6(8), 1066–1081.

Wang, X., et al. (2023). From comfort zone to mortality: Sequence of physiological stress thresholds in Robinia pseudoacacia seedlings during progressive drought. Frontiers in Plant Science, 14. 10.3389/fpls.2023.1149760

Winbourne, J. B., Jones, T. S., Garvey, S. M., Harrison, J. L., Wang, L., Li, D., … & Hutyra, L. R. (2020). Tree transpiration and urban temperatures: Current understanding, implications, and future research directions. BioScience, 70(7), 576–588. 10.1093/biosci/biaa055

Wright, I. J., Reich, P. B., Westoby, M., Ackerly, D. D., Baruch, Z., Bongers, F., … & Villar, R. (2004). The worldwide leaf economics spectrum. Nature, 428(6985), 821–827. 10.1038/nature02403

Xiaoli Fu, Frederick C Meinzer, Metrics and proxies for stringency of regulation of plant water status (iso/anisohydry): a global data set reveals coordination and trade-offs among water transport traits, Tree Physiology, Volume 39, Issue 1, January 2019, Pages 122–134, 10.1093/treephys/tpy087

Xiaoyi Xing, Yarong Jiang, Song Li, Lin Yang, Li Zhang, Wenli Zhu, Research progress in the climate change vulnerability of urban forests, Forestry: An International Journal of Forest Research, Volume 98, Issue 3, July 2025, Pages 309–322, 10.1093/forestry/cpae050

Xu, C., Rahman, M., Haase, D., Wu, Y., Su, M., Pauleit, S., 2020. Surface runoff in urban areas: the role of residential cover and urban growth form. J. Clean. Prod. 262, 121421. 10.1016/j.jclepro.2020.121421.

Yan, F., Sun, Y., Song, F., & Liu, F. (2012). Differential responses of stomatal morphology to partial root-zone drying and deficit irrigation in potato leaves under varied nitrogen rates. Scientia Horticulturae, 145, 76–83. 10.1016/j.scienta.2012.07.026

Zhao, J., Zhao, X., Wu, D., Meili, N., & Fatichi, S. (2023). Satellite-based evidence highlights a considerable increase of urban tree cooling benefits from 2000 to 2015. Global Change Biology, 29(12), 3419–3431. 10.1111/gcb.16667

