## Supplementary material for "Intermediate, rather than Isohydric or Anisohydric, Species were More Resilient to an Extreme Heatwave in a Common Garden in Sydney": Table S1. Summary of tree and shrub species planted in the Hawkesbury Urban Forest common-garden trial (Richmond, NSW, Australia), with family, origin

### Supplementary Information

**Table S1.** Summary of tree and shrub species planted in the Hawkesbury Urban Forest common-garden trial (Richmond, NSW, Australia), with family, origin, and leaf habit. Mean survival is reported for each species (n=32 trees per species). Mean height in 2025, plus AGR and RGR height and diameter, calculated from 2021 to 2025, are reported for species with adequate survival (n=4-30 trees per species). Mean crown dieback in January 2026 is reported for each species (n=3-30 trees per species). Ten species marked with an asterisk (*) were excluded from growth analyses due to survival of <4 individuals in May 2025. Species are ordered by survival, and then by height, in 2025, in descending order. Species (n= 30 species) in bold were measured for plant functional traits.

| Species Name | Family | Origin | Leaf Habit | Mean Survival (%) | Mean Height (cm) 2025 | AGR Height (cm/yr) (mean ± SE) | RGR Height (cm yr-1) (mean ± SE) | AGR Diameter (mean ± SE) | RGR Diameter (mean ± SE) | Dieback (%) |
|---|---|---|---|---|---|---|---|---|---|---|
| ***Eucalyptus bancroftii*** | Myrtaceae | Native | Evergreen | 93.80 | 319.90 | 36.58 ± 5.08 | 0.13 ± 0.02 | 18.50 ± 1.86 | 0.32 ± 0.02 | 4.30 |
| ***Callistemon viminalis*** | Myrtaceae | Native | Evergreen | 93.80 | 276.50 | 40.29 ± 3.15 | 0.20 ± 0.01 | 15.42 ± 1.39 | 0.33 ± 0.02 | 9.49 |
| ***Ulmus parvifolia*** | Ulmaceae | Exotic | Deciduous | 90.60 | 174.60 | 23.36 ± 1.94 | 0.18 ± 0.01 | 6.92 ± 1.19 | 0.25 ± 0.03 | 6.12 |
| ***Pyrus nivalis*** | Rosaceae | Exotic | Deciduous | 87.50 | 189.40 | 13.09 ± 4.16 | 0.06 ± 0.03 | 4.53 ± 0.86 | 0.15 ± 0.04 | 61.31 |
| ***Acacia cognata*** | Fabaceae | Native | Deciduous | 84.40 | 289.90 | 34.83 ± 5.59 | 0.15 ± 0.03 | 7.50 ± 1.64 | 0.15 ± 0.04 | 17.54 |
| ***Acacia melanoxylon*** | Fabaceae | Native | Evergreen | 84.40 | 281.10 | 51.01 ± 6.04 | 0.27 ± 0.02 | 18.56 ± 2.24 | 0.41 ± 0.04 | 8.25 |
| ***Quercus dentata*** | Fabaceae | Exotic | Deciduous | 84.40 | 150.70 | 22.80 ± 3.61 | 0.19 ± 0.03 | 8.10 ± 1.46 | 0.27 ± 0.04 | 45.56 |
| ***Fraxinus oxycarpa*** | Oleaceae | Exotic | Deciduous | 78.10 | 205.80 | 19.11 ± 3.95 | 0.11 ± 0.03 | 6.78 ± 1.21 | 0.20 ± 0.05 | 11.92 |
| ***Cupaniopsis anacardioides*** | Sapindaceae | Native | Evergreen | 78.10 | 82.30 | -0.05 ± 2.13 | -0.03 ± 0.02 | 0.25 ± 0.54 | -0.03 ± 0.04 | 14.29 |
| ***Fraxinus griffithii*** | Oleaceae | Exotic | Deciduous | 75.00 | 129.60 | 19.11 ± 2.59 | 0.19 ± 0.04 | 2.84 ± 0.92 | 0.11 ± 0.06 | 8.92 |
| ***Lophostemon confertus*** | Myrtaceae | Native | Evergreen | 71.90 | 238.30 | 41.55 ± 7.17 | 0.22 ± 0.03 | 4.42 ± 1.95 | 0.05 ± 0.05 | 13.00 |
| ***Eucalyptus albens*** | Myrtaceae | Native | Evergreen | 68.80 | 449.70 | 77.45 ± 10.18 | 0.25 ± 0.04 | 23.83 ± 3.10 | 0.40 ± 0.04 | 4.30 |
| *Melaleuca armillaris* | Myrtaceae | Native | Evergreen | 68.80 | 294.50 | 32.48 ± 5.90 | 0.14 ± 0.03 | 19.39 ± 3.57 | 0.25 ± 0.07 | 8.72 |
| ***Liquidambar styraciflua*** | Altingiaceae | Exotic | Deciduous | 68.80 | 251.70 | 44.22 ± 5.65 | 0.27 ± 0.03 | 11.26 ± 1.57 | 0.30 ± 0.05 | 49.06 |
| ***Quercus palustris*** | Fabaceae | Exotic | Deciduous | 68.80 | 160.10 | 21.24 ± 3.76 | 0.18 ± 0.02 | 7.13 ± 1.00 | 0.32 ± 0.05 | 27.08 |
| *Ficus coronata* | Moraceae | Native | Evergreen | 62.50 | 85.50 | 1.65 ± 4.21 | -0.02 ± 0.05 | -1.05 ± 0.82 | -0.13 ± 0.06 | 54.28 |
| ***Acacia implexa*** | Fabaceae | Native | Evergreen | 59.40 | 400.30 | 79.83 ± 13.05 | 0.29 ± 0.03 | 17.75 ± 2.98 | 0.34 ± 0.04 | 19.99 |
| ***Corymbia eximia*** | Myrtaceae | Native | Evergreen | 56.30 | 407.80 | 82.51 ± 12.29 | 0.32 ± 0.05 | 20.89 ± 3.35 | 0.36 ± 0.05 | 8.46 |
| ***Banksia integrifolia*** | Proteaceae | Native | Evergreen | 56.30 | 348.40 | 63.51 ± 6.03 | 0.29 ± 0.03 | 15.55 ± 1.97 | 0.25 ± 0.02 | 11.67 |
| ***Tristaniopsis laurina*** | Myrtaceae | Native | Evergreen | 56.30 | 180.70 | 6.68 ± 7.42 | -0.01 ± 0.06 | 7.74 ± 2.80 | 0.10 ± 0.08 | 24.28 |
| *Banksia ericifolia* | Proteaceae | Native | Evergreen | 53.10 | 311.40 | 60.77 ± 3.83 | 0.34 ± 0.02 | 15.44 ± 3.48 | 0.29 ± 0.06 | 6.36 |

| Species Name | Family | Origin | Leaf Habit | Mean Survival (%) | Mean Height (cm) 2025 | AGR Height (cm/yr) (mean ± SE) | RGR Height (cm yr-1) (mean ± SE) | AGR Diameter (mean ± SE) | RGR Diameter (mean ± SE) | Dieback (%) |
|---|---|---|---|---|---|---|---|---|---|---|
| ***Eucalyptus globulus*** | Myrtaceae | Native | Evergreen | 50.00 | 389.10 | 71.24 ± 7.32 | 0.27 ± 0.03 | 13.31 ± 2.51 | 0.21 ± 0.05 | 66.38 |
| ***Alphitonia excelsa*** | Rhamnaceae | Native | Deciduous | 50.00 | 152.70 | 28.51 ± 3.07 | 0.28 ± 0.03 | 4.47 ± 0.90 | 0.24 ± 0.05 | 22.27 |
| *Syzygium australe* | Myrtaceae | Native | Evergreen | 50.00 | 99.40 | 7.67 ± 1.93 | 0.09 ± 0.02 | 0.01 ± 0.45 | 0.00 ± 0.04 | 20.44 |
| ***Callistemon pallidus*** | Myrtaceae | Native | Evergreen | 46.90 | 140.40 | 18.61 ± 3.70 | 0.16 ± 0.02 | 2.75 ± 0.69 | 0.16 ± 0.03 | 31.72 |
| ***Eucalyptus rubida*** | Myrtaceae | Native | Evergreen | 40.60 | 419.80 | 78.32 ± 14.97 | 0.27 ± 0.03 | 20.59 ± 3.05 | 0.33 ± 0.04 | 18.73 |
| ***Flindersia australis*** | Rutaceae | Native | Evergreen | 40.60 | 98.90 | 5.87 ± 2.57 | 0.06 ± 0.03 | -0.15 ± 0.64 | -0.04 ± 0.07 | 21.00 |
| *Malus ioensis* | Rosaceae | Exotic | deciduous | 40.60 | 91.20 | 2.81 ± 1.72 | 0.02 ± 0.03 | 3.16 ± 0.53 | 0.18 ± 0.02 | 73.86 |
| ***Magnolia grandiflora*** | Magnoliaceae | Native | Evergreen | 37.50 | 79.50 | -0.52 ± 4.79 | -0.05 ± 0.06 | 2.03 ± 1.74 | -0.02 ± 0.08 | 73.46 |
| ***Syncarpia glomulifera*** | Myrtaceae | Native | Evergreen | 34.40 | 133.20 | 21.81 ± 5.50 | 0.20 ± 0.04 | 1.65 ± 0.63 | 0.09 ± 0.04 | 13.31 |
| ***Acacia longifolia*** | Fabaceae | Native | Evergreen | 31.30 | 330.10 | 57.10 ± 6.89 | 0.27 ± 0.03 | 11.46 ± 2.77 | 0.27 ± 0.05 | 31.14 |
| ***Elaeocarpus reticulatus*** | Elaeocarpaceae | Native | Evergreen | 28.10 | 128.80 | 16.06 ± 18.74 | 0.03 ± 0.10 | 8.35 ± 5.63 | 0.12 ± 0.14 | 52.94 |
| ***Syzygium luehmannii*** | Myrtaceae | Native | Evergreen | 28.10 | 84.00 | 0.86 ± 4.87 | -0.02 ± 0.06 | 13.77 ± 12.14 | 0.19 ± 0.14 | 45.19 |
| ***Eucalyptus mannifera*** | Myrtaceae | Native | Evergreen | 25.00 | 254.80 | 33.88 ± 11.53 | 0.16 ± 0.05 | 11.92 ± 2.78 | 0.24 ± 0.03 | 25.63 |
| *Eucalyptus pauciflora* | Myrtaceae | Native | Evergreen | 21.90 | 340.70 | 36.51 ± 11.40 | 0.15 ± 0.04 | 12.98 ± 4.69 | 0.25 ± 0.08 | 6.67 |
| *Banksia serrata* | Proteaceae | Native | Evergreen | 15.60 | 249.00 | 47.04 ± 5.25 | 0.27 ± 0.03 | 19.40 ± 3.47 | 0.34 ± 0.05 | 13.33 |
| *Banksia marginata* | Proteaceae | Native | Evergreen | 12.50 | 192.50 | 24.57 ± 13.51 | 0.14 ± 0.11 | 7.83 ± 4.67 | 0.03 ± 0.23 | 9.00 |
| ***Quercus ilex*** | Fagaceae | Exotic | Deciduous | 12.50 | 107.00 | 7.50 ± 6.79 | 0.07 ± 0.07 | 3.66 ± 1.65 | 0.22 ± 0.09 | 9.60 |
| *Hymenosporum flavum | Pittosporaceae | Native | Evergreen | 9.40 | 196.66 | NA | NA | NA | NA | 12.50 |
| *Acer palmatum | Sapindaceae | Exotic | Deciduous | 9.40 | 147.66 | NA | NA | NA | NA | 95.17 |
| *Agonis flexuosa | Myrtaceae | Native | Evergreen | 9.40 | 123.60 | NA | NA | NA | NA | 14.25 |
| *Plumeria rubra | Apocynaceae | Exotic | Deciduous | 6.30 | 255.00 | NA | NA | NA | NA | NA |
| *Acacia pendula | Fabaceae | Native | Evergreen | 6.30 | 165.00 | NA | NA | NA | NA | NA |
| *Ficus microcarpa hillii | Moraceae | Native | Evergreen | 6.30 | 60.55 | NA | NA | NA | NA | NA |
| *Ceratopetalum gummiferum | Cunoniaceae | Native | Evergreen | 3.10 | 430.00 | NA | NA | NA | NA | NA |
| *Tibouchina granulosa | Melastomataceae | Exotic | Evergreen | 3.10 | 17.00 | NA | NA | NA | NA | NA |
| *Pipturus argenteus | Urticaceae | Native | Deciduous | 0.00 | NA | NA | NA | NA | NA | NA |
| *Xanthostemon chrysanthus | Myrtaceae | Native | Evergreen | 0.00 | NA | NA | NA | NA | NA | NA |

**Table S2**. Total rainfall and maximum and minimum temperatures across the study years, recorded at the nearest Bureau of Meteorology station (BoM station 067105; Australian Bureau of Meteorology, 2025), located 3.5 km from the study site in Richmond, NSW, Australia. Rainfall data for October and November 2024 were missing but obtained from another station (UWS Hawkesbury, BoM station 067021) located <1 km from our study site.

| Parameters | Year | Jan | Feb | Mar | Apr | May | Jun | Jul | Aug | Sep | Oct | Nov | Dec | Annual |
|---|---|---|---|---|---|---|---|---|---|---|---|---|---|---|
| Rainfall (mm) | 2020 | 40.4 | 280.4 | 102.6 | 20.8 | 27.6 | 17.6 | 103.6 | 45.6 | 23.0 | 96.8 | 54.8 | 122.2 | 935.4 |
| | 2021 | 46.4 | 100.0 | 349.6 | 22.0 | 53.6 | 33.0 | 20.8 | 51.4 | 13.6 | 32.6 | 254.6 | 75.4 | 1053.0 |
| | 2022 | 124.6 | 124.8 | 591.2 | 107.0 | 55.0 | 2.4 | 207.6 | 24.6 | 87.4 | 117.8 | 40.0 | 15.6 | 1498.0 |
| | 2023 | 114.0 | 98.4 | 55.6 | 72.2 | 11.0 | 16.0 | 7.2 | 36.8 | 8.4 | 18.2 | 88.6 | 5.6 | 532.0 |
| | 2024 | 149.2 | 95.0 | 21.0 | 221.0 | 49.6 | 92.8 | 33.2 | 15.4 | 39.6 | 25.8 | 61.8 | 24.6 | 829.0 |
| | 2025 | 37.2 | 52.0 | 85.6 | 82.2 | 269.2 | 3.2 | 69.0 | 167.2 | 55.2 | 12.4 | 47.4 | 57.0 | 937.6 |
| Maximum temperature (°C) | 2020 | 32.5 | 28.9 | 25.8 | 24.5 | 20.3 | 18.6 | 18.2 | 19.2 | 23.1 | 25.7 | 28.3 | 27.1 | 24.4 |
| | 2021 | 29.2 | 27.5 | 25.7 | 24.3 | 20.7 | 17.6 | 18.0 | 20.6 | 23.2 | 25.4 | 23.1 | 26.9 | 23.5 |
| | 2022 | 28.7 | 27.8 | 25.2 | 24.0 | 20.7 | 18.4 | 16.9 | 20.1 | 20.8 | 23.0 | 25.5 | 27.8 | 23.2 |
| | 2023 | 29.2 | 29.9 | 29.0 | 23.5 | 20.3 | 18.9 | 20.2 | 21.3 | 26.0 | 28.0 | 28.1 | 31.3 | 25.5 |
| | 2024 | 29.7 | 29.7 | 28.5 | 24.6 | 20.7 | 17.7 | 17.9 | 21.7 | 24.2 | 25.4 | 28.9 | 32.6 | 25.1 |
| | 2025 | 30.5 | 30.0 | 28.6 | 26.2 | 21.0 | 18.2 | 18.2 | 18.6 | 24.3 | 28.5 | 29.6 | 31.1 | 25.4 |
| Minimum Temperature (°C) | 2020 | 19.8 | 19.1 | 15.5 | 11.8 | 7.2 | 5.3 | 5.1 | 5.3 | 9.4 | 13.0 | 14.6 | 16.5 | 11.9 |
| | 2021 | 17.0 | 17.4 | 15.9 | 10.1 | 8.1 | 5.0 | 4.0 | 4.0 | 6.6 | 10.1 | 14.0 | 15.4 | 10.6 |
| | 2022 | 18.8 | 17.3 | 17.4 | 14.0 | 9.4 | 3.4 | 6.5 | 6.4 | 9.6 | 12.9 | 12.3 | 14.0 | 11.8 |
| | 2023 | 17.5 | 17.4 | 16.4 | 12.0 | 4.4 | 5.2 | 3.3 | 5.4 | 7.3 | 10.3 | 15.6 | 18.2 | 11.1 |

| | | | | | | | | | | | | | |
|---|---|---|---|---|---|---|---|---|---|---|---|---|---|
| 2024 | 19.1 | 19.0 | 16.3 | 12.6 | 9.5 | 5.3 | 5.7 | 8.6 | 7.8 | 11.8 | 16.4 | 17.7 | 12.5 |
| 2025 | 18.0 | 17.8 | 16.0 | 11.8 | 7.6 | 5.1 | 3.8 | 4.7 | 8.0 | 11.3 | 14.2 | 16.2 | 11.2 |

**Table S3**: Results from linear regressions between variables and mean survival in May 2025 for 30 tree or shrub species at an urban site in Richmond, NSW, Australia. Slope, intercept, coefficient of variation ($R^2$), and p-value is provided. Traits are ordered from high to low $R^2$.

| **Trait** | **Slope** | **Intercept** | **$R^2$** | ***p* value** |
|---|---|---|---|---|
| Wood density (g $cm^{-3}$) | 0.001 | 0.521 | 0.090 | 0.107 |
| $\Delta\Psi$ (MPa) | -0.005 | -1.077 | 0.088 | 0.111 |
| Crown dieback (%) | -0.226 | 38.720 | 0.073 | 0.149 |
| Leaf temperature (°C) | 0.017 | 40.430 | 0.066 | 0.170 |
| Relative water content (%) | -0.060 | 53.150 | 0.052 | 0.224 |
| Height (cm) | 0.929 | 174.800 | 0.035 | 0.320 |
| $\Psi_{PD}$ (MPa) | 0.007 | -2.010 | 0.029 | 0.367 |
| Leaf size ($mm^2$) | 17.830 | 1103.000 | 0.029 | 0.371 |
| Stem collar diameter (mm) | 0.177 | 42.390 | 0.021 | 0.448 |
| LMA ($cm^2$ $g^{-1}$) | -0.273 | 105.400 | 0.017 | 0.488 |
| Leaf thickness (mm) | -0.001 | 0.380 | 0.014 | 0.539 |
| $\pi_{tlp}$ (MPa) | -0.001 | -1.792 | 0.013 | 0.555 |
| Leaf width (cm) | 0.001 | 0.212 | 0.012 | 0.561 |
| Thermal safety margin (°C) | -0.013 | -5.045 | 0.005 | 0.706 |
| $g_s$ (mol $m^{-2}$ $s^{-1}$) | 0.000 | 0.043 | 0.004 | 0.736 |
| $\Psi_{MD}$ (MPa) | 0.002 | -3.086 | 0.001 | 0.841 |
| $\Phi_{PSII}$ | -0.000 | 0.280 | 0.001 | 0.887 |
| Stomatal density ($mm^2$) | -0.019 | 68.140 | 0.000 | 0.946 |
| $T_{leafmax}$ (°C) | -0.001 | 47.300 | 0.000 | 0.951 |

**Table S4**: Results from linear or nonlinear regressions between variables and mean crown dieback in January 2026 for 30 tree or shrub species at an urban site in Richmond, NSW, Australia. Model, slope, intercept, coefficient of variation ($R^2$), and p-value is provided. Bold denotes significance at $p \leq 0.05$. Traits are ordered from high to low $R^2$.

| Trait | Model | Slope | Intercept | $R^2$ | *P* value |
|---|---|---|---|---|---|
| $\Delta\Psi$ (MPa) | Quadratic | 167.550 | 136.186 | 0.318 | **0.001** |
| $\Psi_{PD}$ (MPa) | Quadratic | -57.537 | -17.046 | 0.297 | **0.002** |
| Relative water content (%) | Linear | 1.603 | -54.201 | 0.256 | **0.004** |
| Wood density (g $cm^{-3}$) | Linear | -92.234 | 80.098 | 0.207 | **0.012** |
| Leaf width (cm) | Linear | 40.393 | 14.238 | 0.203 | **0.012** |
| Leaf size ($mm^2$) | Linear | 0.003 | 18.353 | 0.163 | **0.027** |
| $\pi_{tlp}$ (MPa) | Linear | -25.237 | -21.602 | 0.089 | 0.109 |
| Survival (%) | Linear | -0.226 | 38.718 | 0.073 | 0.149 |
| Height (cm) | Linear | -0.044 | 35.410 | 0.067 | 0.166 |
| Stem collar diameter (mm) | Linear | -0.152 | 33.346 | 0.050 | 0.233 |
| LMA (g $m^{-2}$) | Linear | 0.069 | 19.048 | 0.030 | 0.361 |
| Leaf temperature (°C) | Linear | 1.951 | -55.616 | 0.025 | 0.409 |
| Thermal safety margin (°C) | Linear | -0.680 | 21.308 | 0.023 | 0.422 |
| Stomatal density ($mm^2$) | Linear | 0.068 | 20.674 | 0.015 | 0.514 |
| $\Psi_{MD}$ (MPa) | Linear | -1.898 | 19.624 | 0.011 | 0.586 |
| $\Phi_{PSII}$ | Linear | 11.214 | 22.219 | 0.003 | 0.773 |
| $T_{leafmax}$ (°C) | Linear | -0.148 | 66.379 | 0.000 | 0.951 |
| $g_s$ (mol $m^{-2}$ $s^{-1}$) | Linear | 11.999 | 24.690 | 0.001 | 0.901 |
| Leaf thickness (mm) | Linear | 3.839 | 23.957 | 0.000 | 0.907 |

**Table S5.** Mean survival and cumulative mortality rates of 48 tree species planted at the common garden in western Sydney, Australia, measured at four time points over a 4.5-year period (2021–2025). Number of seedlings alive and dead, relative to the total of 1536 seedlings, are also provided.

| Year | Month | Alive | Dead | Survival (%) | Mortality (%) | Mortality change (%) | SE (±) |
|---|---|---|---|---|---|---|---|
| 2020 | December | 1536 | 0 | 100 | 0 | 0 | |
| 2021 | December | 1113 | 423 | 72.5 | 27.5 | 27.5 | ±1.140 |
| 2022 | December | 984 | 552 | 64.1 | 35.9 | 8.4 | ±1.224 |
| 2024 | April | 769 | 766 | 50.1 | 49.9 | 14 | ±1.276 |
| 2025 | May | 663 | 872 | 43.2 | 56.8 | 6.9 | ±1.264 |
